# VIPER-TACs leverage viral E3 ligases for disease-specific targeted protein degradation

**DOI:** 10.1101/2024.08.13.607762

**Authors:** Kyle Mangano, Robert G. Guenette, Spencer Hill, Shiqian Li, Seung Wook Yang, Kate S. Ashton, Patrick Ryan Potts

## Abstract

In targeted protein degradation (TPD) a protein of interest is degraded by chemically induced proximity to an E3 ubiquitin ligase. One limitation of using TPD therapeutically is that most E3 ligases have broad tissue expression, which can contribute to toxicity via target degradation in healthy cells. Many pathogenic and oncogenic viruses encode E3 ligases (vE3s), which de facto have strictly limited expression to diseased cells. Here, we provide proof-of-concept for **<u>Vi</u>**ral E3 **<u>P</u>**an-**<u>E</u>**ssential **<u>R</u>**emoving **<u>Ta</u>**rgeting **<u>C</u>**himeras (VIPER-TACs) that are bi-functional molecules that utilize viral E3 ubiquitin ligases to selectively degrade pan-essential proteins and eliminate diseased cells. We find that the human papillomavirus (HPV) ligase E6 can degrade the SARS1 pan-essential target protein in a model of HPV-positive cervical cancer to selectively kill E6 expressing cancer cells. Thus, VIPER-TACs have the capacity to dramatically increase the therapeutic window, alleviate toxicity concerns, and ultimately expand the potential target space for TPD.

## INTRODUCTION

Targeted protein degradation (TPD) is a promising therapeutic approach in which a target protein of interest is degraded by chemically inducing proximity to an effector protein, typically an E3 ubiquitin ligase. Heterobifunctional small molecule (hSM) proteolysis-targeting chimeras (PROTACs) and monovalent molecular glues are the two main chemical modalities used in TPD, which offers several advantages over traditional small molecule inhibitors. First, TPD uses a catalytic rather than an occupancy-based mode of action. Thus, in addition to requiring lower drug concentrations, target ligands that bind non-functional pockets are sufficient to induce degradation^1–3^. Additionally, because protein-protein interactions between targets and E3 ligases contribute to ternary complex formation, ligands can be promiscuous, low affinity and non-inhibitory and yet successful in this application^4–8^.

Currently, small molecule degraders only co-opt a small fraction of the 600+ E3 ligases encoded in the human genome, with the vast majority utilizing either VHL or CRBN^9,10^. Both VHL and CRBN are broadly expressed across cell and tissue types, creating a potential therapeutic liability of target degradation in healthy cells^11–13^. A notable exception which underlines the importance of E3 ligase expression profiles is low VHL expression in platelets. This enables VHL-mediated degradation of BCL-X_L_ to treat certain cancers while avoiding on-mechanism toxicity, thrombocytopenia^14^. Another toxicity challenge with broadly expressed E3 ligases is off-target neo-substrate degradation, a phenomenon reported with CRBN molecular glues^15,16^. These examples highlight the promising opportunity of using tissue-specific or disease-specific E3 ligases in TPD to improve therapeutic windows with degrader modalities.

Like pharmacological TPD, many viruses encode proteins that modulate their hosts’ ubiquitin-proteasome systems to degrade proteins involved in innate immunity and cell cycle regulation. Such viral proteins vary greatly in biophysical characteristics and act through different molecular mechanisms, but ultimately assemble into functional E3 ligase complexes and will be referred to herein as viral E3s (vE3s)^17–19^. Some vE3s act as biologic molecular glues, such as the human papillomavirus (HPV) protein E6, which binds to the host ligase E6AP (UBE3A) to create a p53-recruiting surface^20–23^. On the other hand, many of the described vE3s act as orthogonal substrate receptors in Cullin-RING ligases (CRLs), often hijacking the CUL4 adapter protein DDB1^24–26^. Seven human viruses are recognized as oncoviruses, implicated in approximately 12-17% of all human cancers^27,28^. Bonafide vE3s are expressed by multiple oncoviruses, and while the contribution to carcinogenesis isn’t always clear, the HPV genes *E6* and *E7* and the Hepatitis B virus gene *HBx* incorporate into the host genome and function as oncogenes^29–33^.

TPD as an antiviral approach has recently been reported^34–42^, however, in these cases, a human ligase is used to degrade an essential viral protein. Flipping that paradigm, utilizing vE3s to degrade host proteins represents a novel opportunity for truly disease-specific TPD. Such **<u>Vi</u>**ral E3 **<u>P</u>**an-**<u>E</u>**ssential **<u>R</u>**emoving **<u>Ta</u>**rgeting **<u>C</u>**himeras (VIPER-TACs) could be a therapeutic strategy to treat viral infections or cancers caused by oncogenic viruses with excellent safety profiles. Importantly, because VIPER-TACs will not induce target degradation in healthy cells, the possible target space opens to any of the hundreds of pan-essential proteins. Off-target neo-substrate degradation would also be less of a concern when using vE3s for TPD. In the oncology space, VIPER-TACs would offer comparable therapeutic windows to antibody-drug conjugates (ADCs) or RIPTACs, a recently reported hSM modality which utilizes cancer-specific proteins in inhibitory ternary complexes^43^.

Here, we sought to explore the potential of vE3s as effectors in TPD. Non-inhibitory ligands or ‘silent binders’ to human proteins are seldom reported. Additionally, E3 ligase ligand discovery has historically been challenging. To overcome the lack of available ligands to both the target and effort proteins of interest, we use a chemically induced dimerization (CID) system that brings together two separate fusion proteins to assess vE3 capacity for TPD. We assessed 19 vE3 ligases from 12 different viruses to identify the most active vE3s with on mechanisms TPD. We then show E6-mediated degradation of pan-essential proteins and demonstrate selective cytotoxicity of a tool VIPER-TAC in a chemical biology model of HPV-positive cancer, establishing therapeutic proof-of-concept to utilize vE3s for disease-specific TPD.

## RESULTS

### Establishing a chemically induced dimerization system to assess targeted protein degradation

To assess the capacity of multiple E3 ligases to induce degradation of varying targets without requiring ligand discovery, we developed a dual chimera CID system, combining the previously reported dTAG^44^ and AchillesTAG^45^ degron – ligand pairs (Figure 1A). We synthesized FM4, an hSM comprised of the FKBP12^F36V^ ligand AP1867^46^ linked with 4PEG to the MTH1 ligand from CFT-2139^45^ (Figure 1B), to induce proximity between FKBP12^F36V^-target and MTH1-effector chimeras. The chemical synthesis is described in the supplementary information. By including a HiBiT tag on the target, we can monitor protein abundance after FM4 treatment by luminescence. Benefits of this system include the small sizes of FKBP12^F36V^ (12 kDa) and MTH1 (18 kDa) and the ligands’ high affinities of 2.4 nM and 2.1 nM, respectively^44,45^. Additionally, dTAG offers selectivity over endogenous FKBP12 through a bump-hole ligand, while MTH1 inhibition is expendable in normal and cancerous cells^45^.

**Figure 1.**
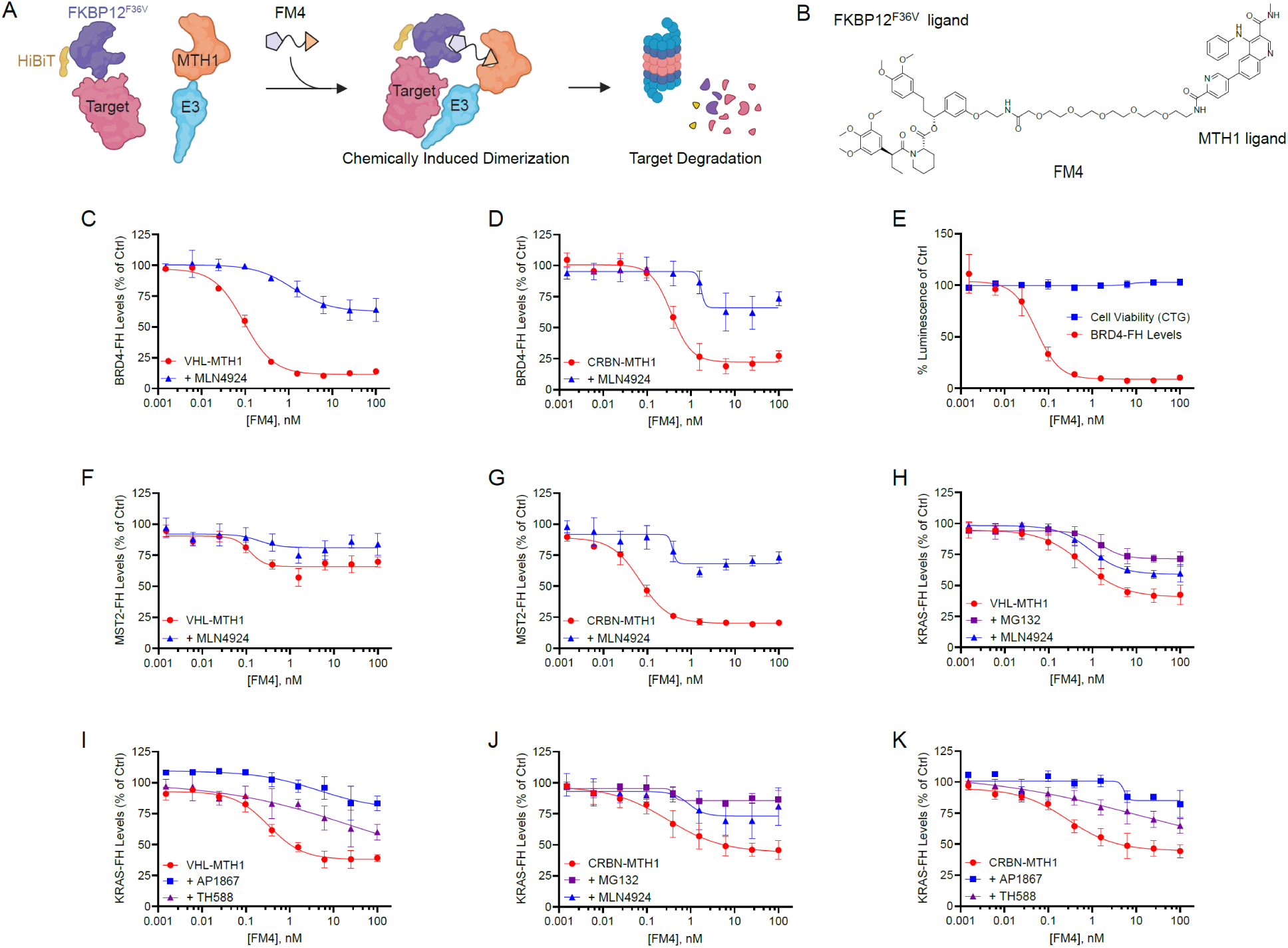
A dual-chimera, chemical induced dimerization system to evaluate E3 ligase capacity for targeted protein degradation. A) Cartoon depiction of the CID system. HiBiT tag on the target allows for measuring abundance. Created with BioRender.com B) Structure of the FM4 heterobifunctional small molecule. C-D) Degradation of HiBiT-BRD4^BD1/2^-FKBP12^F36V^ by VHL-MTH1 and CRBN-MTH1, measured using the Nano-Glo HiBiT Lytic Detection System 24hr after treatment with FM4, relative to a DMSO control. Rescue by 15-minute pre-treatment with 1 μM of the neddylation inhibitor MLN4924. Data representative of three independent experiments, reported as the mean ± S.E. E) Cell viability measured by CellTiterGlo assay 24hr after FM4 treatment, relative to a DMSO control, in cells transiently transfected with VHL-MTH1 and HiBiT-BRD4^BD1/2^-FKBP12. Data representative of two independent experiments reported as the mean ± S.E. F-G) Degradation of HiBiT-MST2-FKBP12^F36V^ by VHL-MTH1 and CRBN-MTH1 24hr after FM4 treatment, with rescue by 15-minute pre-treatment with 1 μM MLN4924. Data representative of three independent experiments, reported as the mean ± S.E. H-K) Degradation of HiBiT-FKBP12^F36V^-KRAS by VHL-MTH1 and CRBN-MTH1 24hr after FM4 treatment. *H* and *J* show rescue of degradation by 15-minute pre-treatment with 10 μM of the proteasome inhibitor MG132 or 1 μM MLN4924. *I* and *K* show rescue of degradation by 15-minute pre-treatment with 5 μM of the FKBP12^F36V^ ligand AP1867 or 2 μM of the MTH1 ligand TH588. Data representative of three independent experiments, reported as the mean ± S.E.

We first validated the system with the well characterized E3 ligases CRBN and VHL paired with HiBiT-BRD4^BD1/2^-FKBP12^F36V^ (Bromodomains 1 and 2) as a target via transient expression in human embryonic kidney 293T (HEK293T) cells. Both VHL-MTH1 (Figure 1C) and CRBN-MTH1 (Figure 1D) degraded BRD4 in a FM4 dose dependent manner, with maximum degradation (D_max_) values of 90% and 81%, respectively, compared to DMSO treated control cells at 24hr. BRD4 degradation is reduced by pre-treatment with the neddylation inhibitor MLN4924 (Figure 1C-D), indicating the MTH1-ligase chimeras are acting on-mechanism in CRL complexes. FM4 is highly potent, with DC_50_ values of 0.09 - 0.22 nM, and no cytotoxicity was observed at 100 nM FM4, the maximum concentration in the assay (Figure 1E). To show broad applicability with various targets, the cytosolic kinase MST2 and membrane associated KRAS were tested for degradation. Interestingly, VHL is a poor degrader of HiBiT-MST2-FKBP12 ^F36V^ (Figure 1F) while CRBN degraded 80% of MST2 (Figure 1G). The VHL target specificity indicates that this system can report on ligase-target compatibility, and the degradation is not driven by the FKBP12 and HiBiT tags.

KRAS was N-terminally FKBP12-tagged (HiBiT-FKBP12^F36V^-KRAS) because its C-terminus is important in membrane localization. VHL degraded 59% of KRAS, with rescue by pre-treatment with the proteasome inhibitor MG132 and partial rescue by MLN4924 (Figure 1H). KRAS degradation requires proximity of the fusion proteins, demonstrated by pre-treatment with an excess of competitive binders to either tag, FKBP12^F36V^ ligand AP1867 or the MTH1 ligand TH588^47^ (Figure 1I). Highly similar degradation profiles were observed with CRBN (Figure 1J-K). Importantly, FM4 did not significantly alter KRAS levels in cells without MTH1-E3 expression (Figure S1A). To probe whether the geometry of substrate recruitment in the CID system matters, we evaluated whether N-terminal or C-terminal MTH-tagged ligases performed differently. Intriguingly, N-terminal MTH1-tagged ligases were far less active than C-terminal MTH-tagged ligases (Figure S1B), suggesting orientation of substrate recruitment is critical for efficient degradation. Taken together, dual chimera, FM CID system can be used to assess E3 ubiquitin ligase ability to induce degradation of target proteins.

### Viral E3 ligases demonstrate capacity to induce neo-substrate degradation

After validating our CID approach with well-characterized human ligases, we asked if vE3s could be redirected to degrade a target of interest (Figure 2A). A literature search identified putative vE3s^17,18,25,48–52^, 19 of which were tagged and assessed for their capacity to degrade BRD4 and MST2 in the FM4 CID system. A summary of the D_max_ from FM4 titration experiments demonstrates that many vE3s have the capacity to induce target degradation (Figure 2B). Example titration curves of select vE3s (E6 from HPV, UL145 from CMV, V from HPIV2, and NSP1 from rotavirus) with rescue by MG132 are shown in Figure 2C-F. About half of the ligases showed a difference between N- and C-terminal MTH1 tag orientations (Figure S2). Most of the vE3s induced at least 40% degradation of BRD4, while only seven degraded at least 40% of MST2. Some vE3s showed strong target selectivity (Figure 2B), while others degraded BRD4 and MST2 to similar extents, such as E6 and UL145 (Figure 2C-D). Activity was seen from vE3s encoded in different types of viruses and that act by various host ligases, both CRL and non-CRL mechanisms (Figure 2B). These results established that many vE3s are competent for neo-substrate degradation and prioritized the most active vE3s to move forward with mechanistic studies.

**Figure 2.**
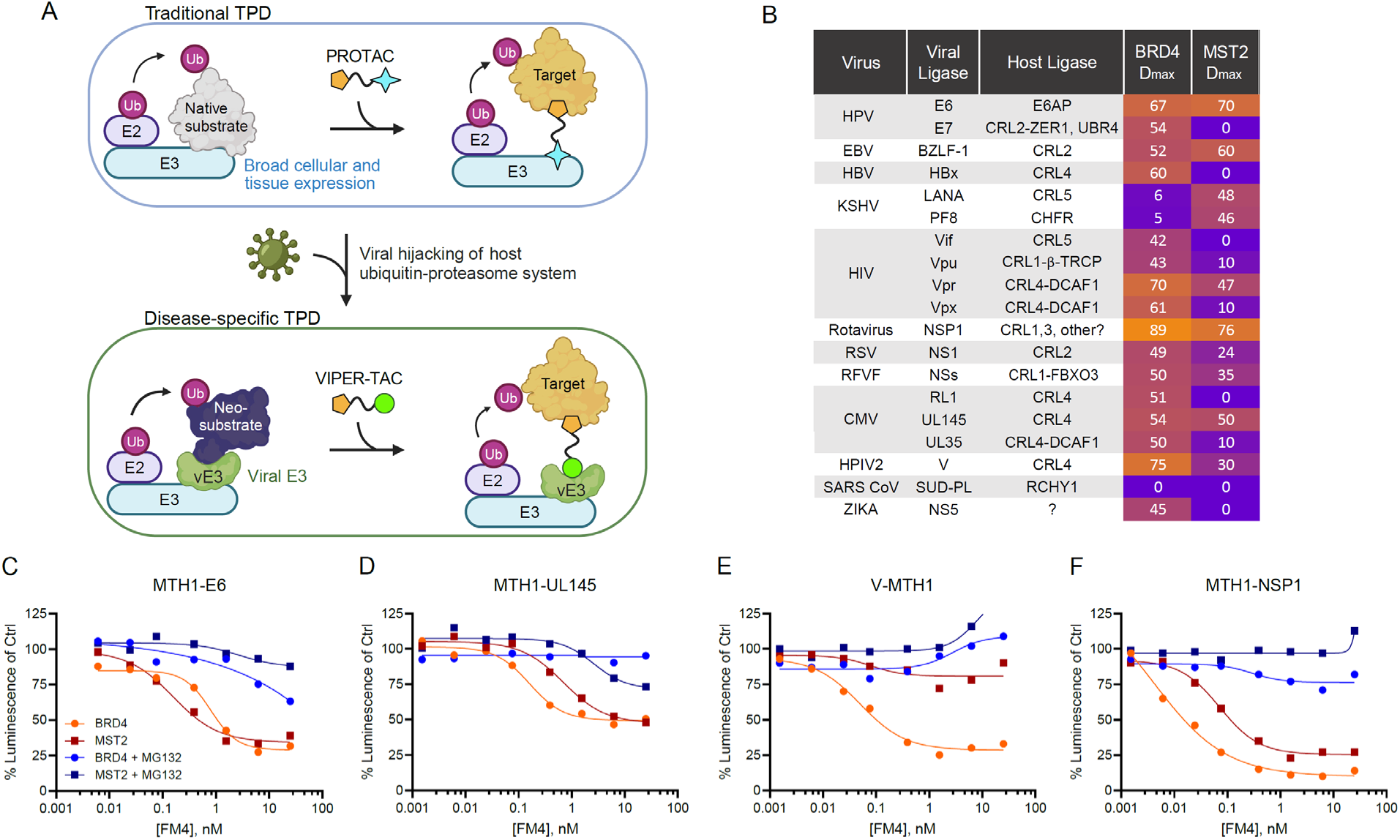
Screening a panel of 19 viral E3 ligases for TPD capacity. A) Cartoon depicting how viruses hijack the host ubiquitin proteasome system, and the concept of chemically redirecting viral E3 ligases (vE3s) for disease-specific TPD. Created with BioRender.com. B) A selection of vE3-MTH1 chimeras was tested in the FM4 CID assay against HiBiT-BRD4^BD^^1/2^-FKBP12^F36V^ and HiBiT-MST2-FKBP12^F36V^. Maximum target degradation (D_max_) from FM4 dose responses, relative to a DMSO control at 24hr are reported. Data representative of one experiment, see also Figure S2 for comparisons of MTH1 tag orientations. C-F) FM4 dose response curves of select viral ligases from the screen in panel *B*. Proteasome-dependent degradation was confirmed by 15-minute pre-treatment with 10 μM MG132. Data representative of one experiment.

### Mechanistic activity of select MTH1-tagged vE3s

Of the vE3s with the strongest degradation profiles, E6, HBx, UL145, V, and Vpr have previously described binding-deficient mutants that disrupt interaction with host ligase components. To assess if the activity in the CID screen was dependent on these definable interactions, mutant and wildtype vE3s were compared for human ligase binding and CID activity. First, we examined whether E6AP binding by E6 is required for CID activity. E6AP co-precipitated with MTH1-E6^WT^ (HPV16 variant was used throughout) but not MTH1-E6^L57E^ or MTH1-E6^6Mut^ which contains six mutations in the E6AP binding groove^21^ (Figure 3A). Importantly, both E6 mutants reduce the ability of E6 to degrade MST2 in the CID assay (Figure 3B). Second, we examined whether V activity is dependent on DDB1 binding. DDB1 co-precipitated with V^WT^-MTH1 but not with V^Δ32^-MTH1 which has a deletion of the N-terminal DDB1 binding H-box helix^24^ (Figure 3C). Importantly, the non-DDB1 binding V^Δ32^-MTH1 failed to degrade BRD4 in the CID assay (Figure 3D). Similarly, DDB1 binding (Figure 3E) and CID activity (Figure 3F) were reduced with MTH1-UL145^5Mut^ which contains five mutations in the H-box alpha helix^25^. On the other hand, DDB1 binding-deficient HBx^24,53^ (Figure S3A) and DCAF1 binding-deficient Vpr^54^ (Figure S3B) did not rescue activity in the CID assay, indicating that target degradation mediated by these MTH1-tagged vE3s may not occur through the predicted mechanism.

**Figure 3.**
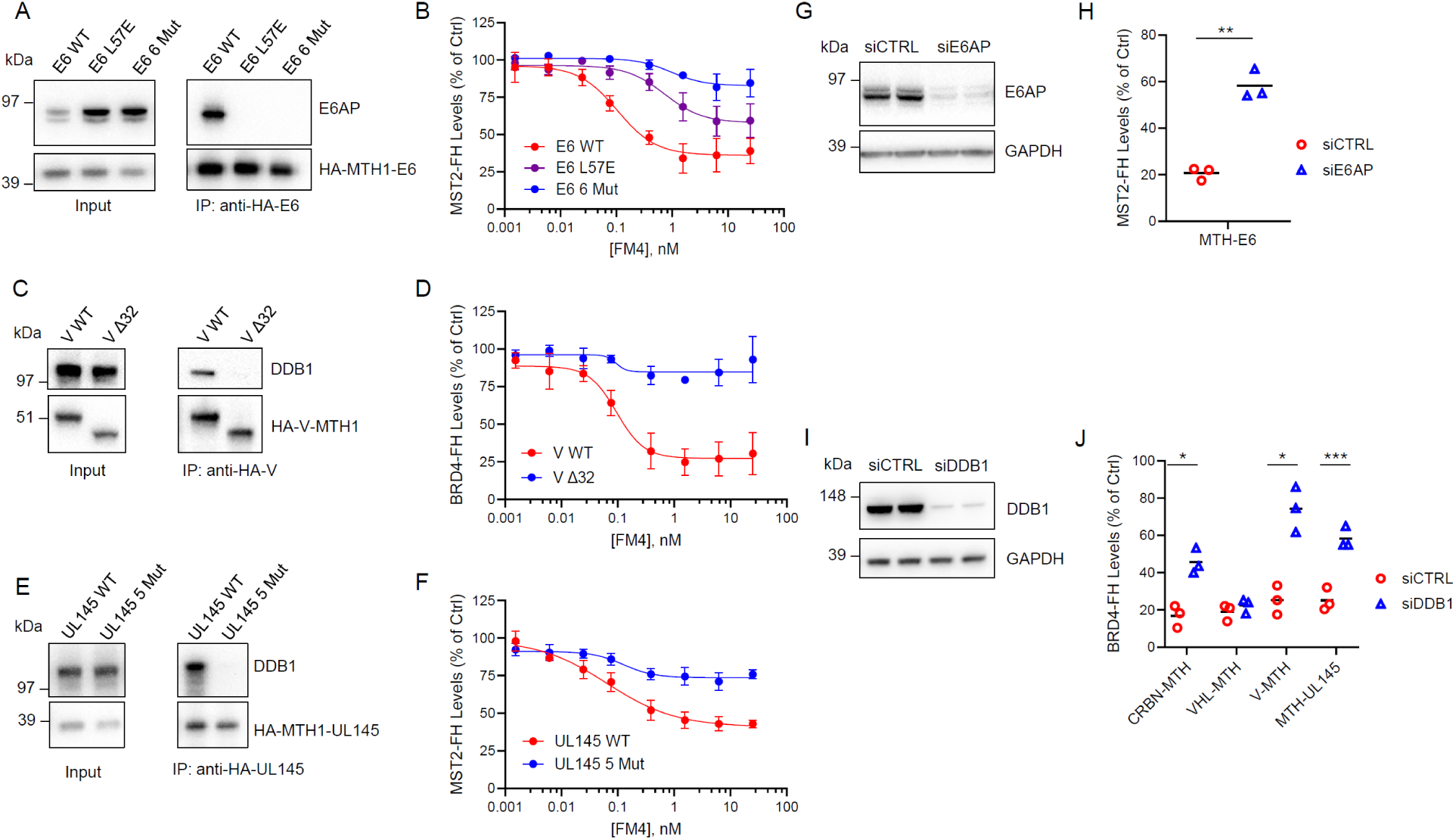
The vE3s E6, V, and UL145 promote on mechanism TPD. A) Co-immunoprecipitation (Co-IP) with anti-HA magnetic beads demonstrates that HA-MTH1-E6 retains the expected binding mode to the human E6AP ligase. The single point mutation L57E or six amino acids substitutions in the E6AP binding pocket (Y39A, L57E, R62A, Y77A, S81A and R138A) reduce E6AP binding compared to wild-type (WT). Representative images shown of two independent experiments for all co-immunoprecipitation experiments. B) E6 mutants exhibit reduced HiBiT-MST2-FKBP12^F36V^degradation in the CID assay after 24hr FM4 treatment. All HiBiT luminescent data representative of three independent experiments, reported as the mean ± S.E. C) Co-IP demonstrates that V-MTH1-HA retains expected binding mode to human CRL4 adapter protein DDB1. V^Δ32^ mutant has the first 32 amino acids truncated that is required for DDB1 binding. D) V^Δ32^ mutant exhibits reduced HiBiT-BRD4^BD1/2^-FKBP12^F36V^ degradation in the CID assay after 24hr FM4 treatment. E) Co-IP demonstrates that HA-MTH1-UL145 retains expected binding mode to DDB1. Five amino acids substitutions in the UL145 H-box motif (N25A, L29A, L30A, R33A and R35A) reduces DDB1 binding. F) UL145 mutant exhibits reduced HiBiT-MST2-FKBP12^F36V^ degradation in CID assay after 24hr FM4 treatment. G) Knockdown efficiency of E6AP at time of HiBiT readout, 72hr post siRNA treatment. Biological replicates are shown. H) HiBiT-MST2-FKBP12^F36V^ levels 24hr after treatment with 6 nM FM4 shows significantly reduced degradation in E6AP knockdown vs control cells (paired t-test, ** = P ≤ 0.01), measured using the Nano-Glo HiBiT Lytic Detection System. Individual data points from three independent experiments are presented, black lines represent the means. I) Knockdown efficiency of DDB1 at time of HiBiT readout, 72hr post siRNA treatment. Biological replicates shown. J) HiBiT-BRD4^BD1/2^-FKBP12^F36V^ levels 24hr after treatment with 6 nM FM4 shows significantly reduced degradation in DDB1 knockdown vs control cells (paired t-test, * = P ≤ 0.05, *** = P ≤ 0.001). Individual data points from three independent experiments are presented, black lines represent the means.

To further validate the mechanism of action, E6AP and DDB1 were knocked down with siRNA 24hr prior to the CID assay. Consistent with the vE3 mutation data, E6AP knockdown (Figure 3G) significantly reduced E6-mediated MST2 degradation (Figure 3H) while DDB1 knockdown (Figure 3I) reduced the ability of V-MTH1 and MTH1-UL145 to degrade BRD4 (Figure 3J). CRBN-MTH1, a CRL4^DDB1^ E3, was included as a positive control and showed dependence on DDB1 level, whereas VHL-MTH1, a CRL2^EloB/C^ E3, was immune to DDB1knockdown (Figure 3J). Consistent with the off-mechanism activity seen with their mutants, degradation with HBx and Vpr was unchanged in DDB1 knockdown (Figure S3C). Overall, MTH1-tagged E6, V, and UL145 act through their literature-described mechanisms, requiring the host ligase E6AP and CRL adapter protein DDB1, respectively, to induce degradation of target proteins in the FM4 CID assay.

### Target selection in cervical cancer cells for E6-mediated degradation

A major advantage to utilizing a disease-specific E3 ligases is that any protein can be degraded for therapeutic effect, including pan-essentials to induce cell death. We chose E6 as the vE3 to demonstrate the VIPER-TAC concept because it is highly active against BRD4 and MST2, demonstrates on mechanism activity, and is a well-characterized viral oncogene affecting a large global patient population^32,55^. Homozygous HiBiT-FKBP12^F36V^ knock-in at an endogenous target locus would enable FM4 cytotoxicity in cells co-expressing MTH1-E6. To select pan-essential genes to screen with E6 in the FM CID system, we ranked genes based on their average Chronos dependency scores across 1100 DepMap cell lines (Figure 4A, DepMap 23Q4). We chose eight of the most essential targets representing a variety of cellular processes, including: protein synthesis and folding, nuclear transport, cell cycle regulation and splicing. FM4 titration showed dose-dependent degradation of multiple essential proteins (Figure 4B) while revealing eIF2S1 and SARS1 as the most susceptible targets, with each degraded over 60%.

**Figure 4.**
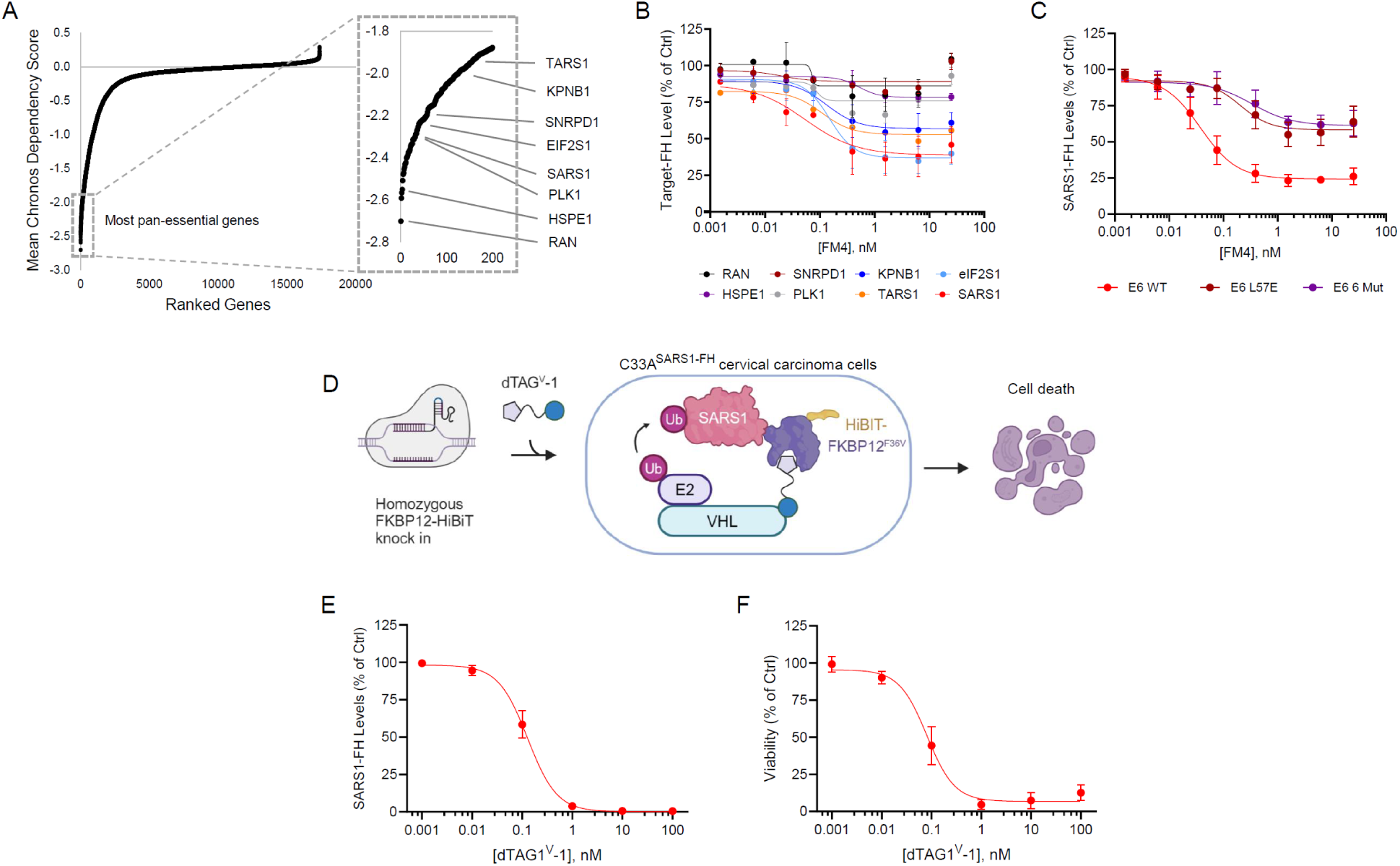
Pan-essential target selection for FKBP12 knock-in in C33A cervical cancer cells. A) Genes ranked by average Chronos dependency score across 1100 cell lines (DepMap 23Q4). Zoomed box shows the top 200 pan-essential genes, with genes selected for further testing marked. B) MTH1-E6 mediated degradation of C-terminally FKBP12^F36V^ tagged HiBiT-pan-essential targets following 24hr FM4 treatment, as described in Figure 1. All data representative of three independent experiments, reported as the mean ± S.E. C) Reduced HiBiT-SARS1-FKBP12^F36V^ degradation with E6 mutants in the CID assay after 24hr FM4 treatment indicates SARS1 degradation is dependent on E6 interaction with E6AP. D) Cartoon illustrating that after homozygous knock-in of FKBP12^F36V^-HiBiT at endogenous loci of a pan essential gene like SARS1, dTAG^V^-1 treatment will lead to cell death if target protein is sufficiently degraded. C33A is a cervical carcinoma cell line, C33A^SARS1-FH^ denotes a clonal cell line with homozygous knock-in of the tag at the SARS1 locus. Created with BioRender.com. E) SARS1-FKBP12^F36V^-HiBiT abundance 4hr post dTAG^V^-1 treatment in C33A^SARS1-FH^, relative to DMSO control. F) Cell viability measured by CellTiterGlo assay 48hr post dTAG^V^-1 treatment in C33A^SARS1-FH^, relative to DMSO control.

While HPV-positive, E6 expressing cancer cell lines such as CASKI, HELA and SIHA are commonly cultured, knocking in the MTH1 tag to create endogenous MTH1-E6 is challenging because the *E6* gene is incorporated at multiple loci^30^. We chose to use C33A cells, an HPV-negative, p53 mutant cervical carcinoma cell line, in which stable E6 expression can be used as a model of HPV-positive cervical cancer^56,57^. SARS1 (seryl-tRNA synthetase 1) is the sixth most essential gene in C33A (Figure S4A, DepMap 23Q4) while ranking 9^th^ in SIHA and 70^th^ in CASKI, all higher rankings than eIF2S1. The E6AP binding-deficient E6 mutants show less activity against SARS1 in the FM4 CID system (Figure 4C) and an increased Dmax of 80% was observed with E6^WT^ in this experiment, leading us to attempt knock-in at the endogenous SARS1 locus in C33A cells.

Using gene editing technology followed by limiting dilution cloning and phenotypic screening (Figure 4D), we generated a clonal C33A line with homozygous FKBP12^F36V^-HiBiT knock-in at the C-terminus of endogenous SARS1 (C33A^SARS1-FH^). Treatment with dTAG^V^-1 caused 99% degradation of SARS1-FH at 4hr (Figure 4E) and killed the cells within 48hr (Figure 4F), indicating an absence of a wildtype SARS1 allele. Genomic PCR (Figure S4B) and immunoblotting with anti-SARS1 antibodies (Figure S4C) confirmed the homozygous knock-in.

### Selective cytotoxicity and mechanism of action of VIPER-TAC treatment in E6 expressing cells

MTH1-E6 was stably expressed at the adeno-associated virus integration site 1 (AAVS1) locus in C33A^SARS1-FH^ driven by either a constitutive or dox-inducible promoter. A nearly complete loss of p53 in the constitutive C33A^SARS1-FH:E6^ cells indicates that MTH1-E6 retains its native functionality (Figure 5A, lane 4), and this cell line was used in subsequent experiments.

**Figure 5.**
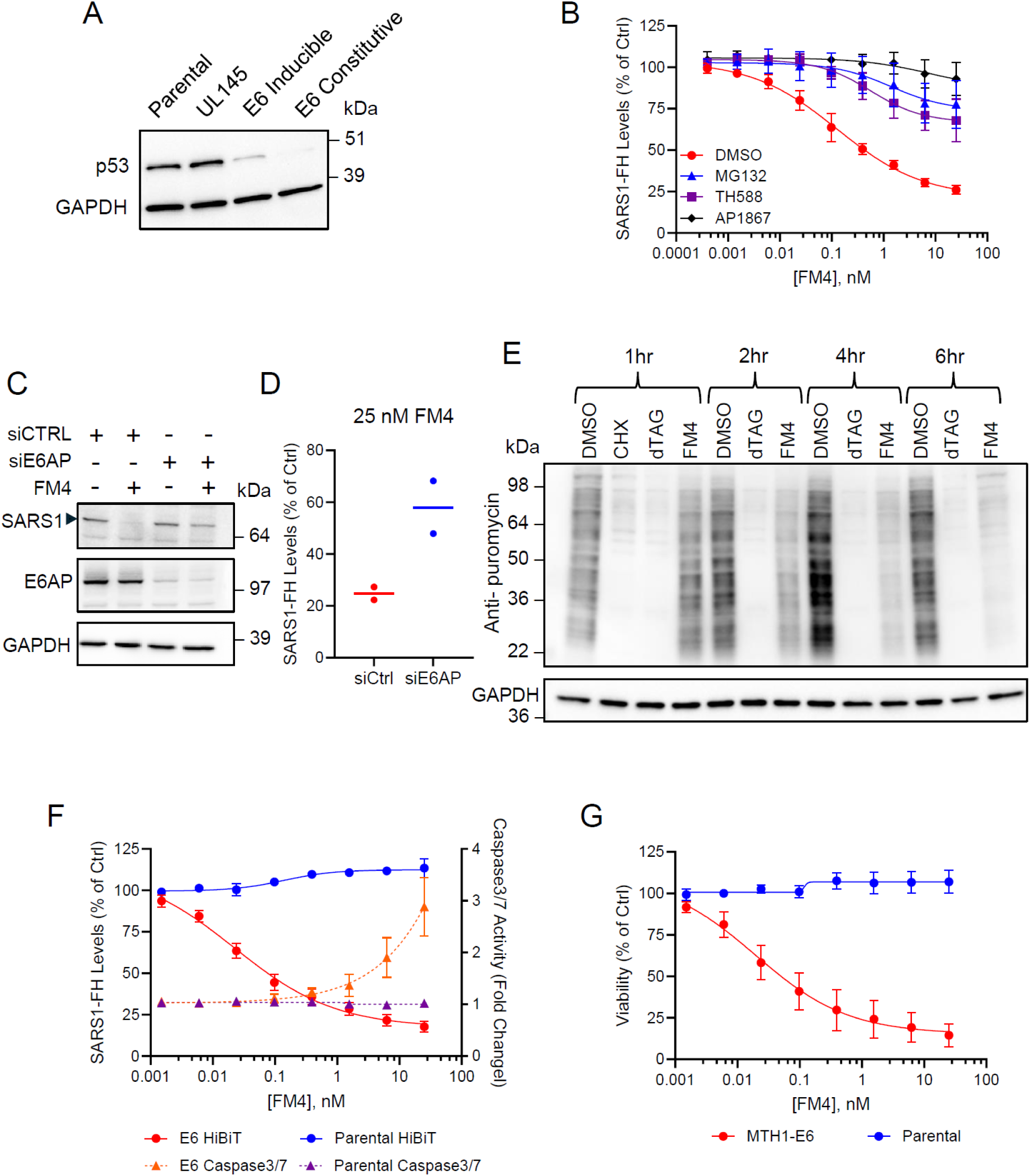
Stable MTH1-E6 expression in C33A SARS1-FKBP12-HiBiT cells enables demonstration of therapeutic proof-of-concept for disease-specific TPD by VIPER-TAC. A) Stable expression of MTH1-E6 from the AAVS1 safe harbor locus in C33A^SARS1-FH^ cells leads to degradation of endogenous p53. E6 pooled line in lane 4 is driven by constitutive promoter and was used in subsequent experiments (C33A^SARS1-FH: E6^), E6 cells in lane 3 is driven by dox-inducible promoter. Representative image shown of two independent experiments. B) SARS1-FKBP12^F36V^-HiBiT degradation after 6hr FM4 (a VIPER-TAC in this context) treatment in C33A^SARS1-FH: E6^ cells, relative to DMSO control, measured using the Nano-Glo HiBiT Lytic Detection System. Rescue by pre-treatment with 10 μM MG132, 5 μM AP1867 and 2 μM TH588 indicate that the degradation is proteasome and proximity dependent. Data representative of three independent experiments, reported as the mean ± S.E. C) Knockdown of E6AP by siRNA prevents FM4-induced degradation of SARS1-FKBP12^F36V^-HiBiT in C33A^SARS1-FH: E6^ cells. Cells treated with 25 nM FM4 for 6hr before lysis and Western blotting with anti-SARS1 antibody. Representative images shown of two independent experiments. D) SARS1-FKBP12^F36V^-HiBiT degradation after same siRNA and FM4 treatment conditions as panel *C*, measured by HiBiT luminescence using the Nano-Glo HiBiT Lytic Detection System. Individual data points from two independent experiments are presented, lines represent the means. E) Measurement of puromycin incorporation into nascent proteins shows that FM4 treatment globally reduces protein synthesis within 2 hr. C33A^SARS1-FH; E6^ cells were treated with 30 μg/mL (w/v) cycloheximide (CHX), 10 nM dTAG^V^-1, or 25 nM FM4 for 1 to 6 hr and then pulse-labeled with 50 μg/mL (w/v) puromycin for 30 minutes before analysis by Western blotting with an anti-puromycin antibody. Representative images shown of two independent experiments. F) Caspase-3/7 activity was measured 6 hr after treatment with 25 nM FM4 in C33A^SARS1-FH: E6^ cells by the Caspase-Glo 3/7 assay following the manufacturers protocol. Data representative of three independent experiments, reported as the mean ± S.E. G) FM4 VIPER-TAC treatment is selectively cytotoxic in C33A^SARS-FH: E6^ cells compared to the parental C33A^SARS-FH^ cells in a 48hr CellTiterGlo assay. Data representative of three independent experiments, reported as the mean ± S.E.

Excitingly, a 6hr treatment with FM4 (a proof-of-concept VIPER-TAC in this cell line) causes proteasome- and proximity-dependent degradation of SARS1 in the C33A^SARS1-FH:E6^ cells, as indicated by rescue with MG132 or competitive FKBP12 ligand AP1867 and MTH1 ligand TH588 (Figure 5B). Knockdown of E6AP with siRNA 24hr prior to 25 nM FM4 treatment reduced degradation from 75% to 35% by western blotting (Figure 5C) and luminescence (Figure 5D). Degradation of SARS1, an aminoacyl tRNA synthetase, should decrease the pool of charged serine and selenocysteine tRNAs in the cell, thus inhibiting global protein synthesis as ribosomes stall at the corresponding mRNA codons. Puromycin labeling^58^ was performed to measure protein synthesis in the C33A^SARS-FH:E6^ cells. A time course of FM4 treatment reveals greatly reduced protein synthesis after 4hr (Figure 5E). Inhibition of protein synthesis can be cytostatic, but after just 6hr FM4 treatment, Caspase3/7 activity is detectable in the MTH1-E6 expressing cells, indicating TPD-induced apoptosis (Figure 5F).

Critically, FM4 is selectively cytotoxic in the MTH1-E6 expressing cells compared to the parental cell line (Figure 5G). At 48hr post FM4 treatment, there is an 85% reduction in cell viability via Cell TiterGlo assay, with an EC_50_ of 0.2 nM. FM4 treatment has no effect on cell viability in the C33A^SARS1-FH^ parental cell line without MTH1-E6, demonstrating the VIPER-TAC therapeutic proof-of-concept. Overall, we demonstrate rapid, on-mechanism degradation of the pan-essential SARS1 protein by VIPER-TAC treatment only in cancer cells expressing the vE3 ligase E6, resulting in reduced protein synthesis and induction of apoptosis.

## DISCUSSION

Here, we have described a proof-of-concept study showing that viral E3 ubiquitin ligases can be utilized by VIPER-TACs for disease-specific TPD. A chemically induced dimerization system was developed in which an hSM, FM4, brings together two separately tagged proteins. We initially tested the system with a panel of viral E3 ubiquitin ligases, but it could be readily adapted to evaluate induced proximity outcomes with different classes of effector proteins. After identifying the pan-essential protein SARS1 as a degradable target of the papillomavirus protein E6, we genetically engineered a relevant cancer cell line to be reliant on homozygous FKBP12-tagged SARS1. FM4 treatment is benign in that parental cell line but becomes potently cytotoxic in cells stably co-expressing MTH1-E6, thus demonstrating the potential of the VIPER-TAC modality for selective and potent therapeutic effects while maintaining large therapeutic windows. This approach dramatically widens the target space to include many proteins that have been traditionally off the table due to toxicity concerns.

Although the HPV vaccines introduced almost two decades ago effectively prevent infection and thus HPV-positive cancers^59^, there remains a large unmet medical need. High-risk HPV subtypes 16 and 18 are estimated to contribute to 500,000 new cancer cases per year, the majority of which being cervical carcinoma^55,60^. Therapeutically targeting E6 has been explored with a variety of modalities, including attempts at disrupting the E6-E6AP interaction with small molecules and peptides^61–63^. However, the large protein-protein interface and picomolar interaction make in-cell disruption of the E6-E6AP complex very challenging^23^. We demonstrated an alternative approach which redirects the intact E6-E6AP complex toward degradation of an essential human protein. A small molecule degrader utilizing endogenous E6 could compete with clinical stage E6/E7 targeted T-cell therapies^64^ or arenaviruses^65^ while potentially offering oral dosage forms, lower manufacturing costs and easier distribution.

Since most of the vE3s showed some activity in the CID screen, we believe this therapeutic approach could be expanded to treat other vE3 expressing cancers, as well as viral infections. Additionally, TPD-based antimicrobials can be expanded to utilize other pathogen E3 ubiquitin ligases. This has been recently demonstrated in bacteria^66,67^ and could be especially useful in combating eukaryotic infectious agents, which are often difficult to treat due to close homology of traditionally druggable targets. As for human disease-specific E3 ligases, some members of the melanoma antigen (MAGE) gene family associate with E3s and although normally restricted to the germ line, are aberrantly expressed in certain cancers^68^. Whether or not MAGEs can be repurposed for TPD remains to be seen. Utilizing cell type- and tissue-specific ligases is another promising avenue for increasing therapeutic windows with degrader modalities^13^.

In conclusion, our work demonstrates that viral E3 ubiquitin ligases have the capacity for use in targeted protein degradation. Future work should focus on ligand discovery to vE3s and other disease-specific E3 ligases and finding optimal target proteins for such cases when the therapeutic selectivity is effector-mediated. Exploring other types of disease-specific effector proteins for use in chemically induced proximity may also yield new therapeutic hypotheses.

## ACKNOWLEDGEMENTS

We thank Jonathan W. Bushman, Dhanusha A. Nalawansha, Christopher E. Smith, Marco Paolo Jacinto, Xiaorui Fan, Han-Yuan Li, and all members of IPP and the Amgen postdoc community for helpful discussions and comments on the manuscript.

## AUTHOR CONTRIBUTIONS

Conceptualization, K.M. and P.R.P.; Methodology, K.M., R.G.G., S.H., S.L., S.W.Y., K.S.A., and P.R.P.; Investigation, K.M., R.G.G., S.H., and K.S.A.; Writing – Original Draft, K.M., P.R.P.; Writing – Review and Editing, K.M., R.G.G., S.W.Y., and P.R.P.; Supervision, S.W.Y. and P.R.P.

## METHODS

### Cell lines and reagents

Human embryonic kidney 293T (HEK293T) cells were obtained from ATCC (CRL-3216) and cultured in DMEM (Gibco) supplemented with 10% (v/v) heat inactivated fetal bovine serum (FBS) (Gibco). C33A cells were obtained from ATCC (HTB-31) and cultured in RPMI 1640 Medium (Gibco) supplemented with 10% (v/v) heat inactivated FBS (Gibco). All media were sterile filtered prior to usage (Nalgene #564-0020) and cultured with 1 µg/mL (w/v) puromycin (Thermo Scientific J67236-XF) and 10 µg/mL (w/v) blasticidin S (Thermo Scientific J67216-XF) as needed. Cells were cultured at 37 °C and 5% CO_2_ in a humidified incubator and regularly checked for mycoplasma infection. Plasmid transfection was performed using Lipofectamine LTX with PLUS Reagent (Invitrogen 15338-100), and siRNA transfection was performed using Lipofectamine RNAiMAX (Invitrogen 13778075). Compounds were dissolved in DMSO and all treatments were performed in complete medium. MG132 (474791) and MLN4924 (50-547-70001) were purchased from Millipore Sigma. AP1867 (HY-114434) was purchased from MedChemExpress. TH588 (5334) and dTAG^V^-1 (6914) were purchased from Tocris. M3814 (S8586) was purchased from Selleckchem.

### Plasmids and cloning

The pcDNA3.1(+) vector (Invitrogen V79020) was used to express MTH1 tagged E3 ligases and HiBiT-FKBP12 tagged targets with GSG or GSGSG linkers between dimerization tags and target or E3 ligase proteins. Oligonucleotides and synthetic genes were obtained from IDT and Genscript. Cloning was performed using In-Fusion Snap Assembly (Takara 638948). Plasmid amplifications were performed in Stellar Competent Cells (Takara 636763). Plasmids were confirmed with Sanger sequencing. To generate the C33A^SARS1-FH^ cells (FKBP12-HiBiT-P2A-BSD knock-in at the SARS1 endogenous genomic locus at C-terminus in C33A), four plasmids were used, obtained from Genscript or VectorBuilder. pKM1 contains the insertion sequence plus homology directed repair arms in the pUC57-Mini vector, pKM2 contains codon optimized editing enzyme in the pCAG vector, pKM3 is sgSARS1 (5’-AGGCAGGAATGTTCAAGCAT -3’) and a puromycin resistance gene in the pUC57 vector, pKM4 contains the i53 (inhibitor of 53BP1) coding sequence with EF1a promoter in the pUC57 vector. To express MTH1 tagged ligases at the AAVS1 locus in C33A^SARS1-FH^ cells, two plasmids were used. pKM5 contains editing enzyme and AAVS1 sgAAVS1 (5’-GCTAGTGGCCCCACTGTGGGGG -3’) in pUC57 vector. pKM6 contains the expression sequences and puromycin selection marker with AAVS1 homology directed repair arms in pUC57 vector.

### Cell line engineering

To generate the C33A^SARS1-FH^ cells (FKBP12-HiBiT-P2A-BSD knock-in at the SARS1 endogenous genomic locus in C33A), C33A cells were seeded overnight in 12 well plates at 6×10^6^ cells per well. The following day, 0.4 μg pKM1, 0.3 μg pKM2, 0.2 μg pKM4 and 0.1 μg pKM3 were transfected with 1 μL PLUS reagent and 3 μL Lipofectamine LTX, delivered in 100 μL Opti-MEM (Gibco). 6 hours after transfection, cells were passaged 1:6 into 6 well plates with 2.5 mL tissue culture medium and 1 μM M3814 and incubated for 24 hours. The media was replaced with media containing 1 μg/mL (w/v) puromycin and 1 μM M3814 and cells were incubated for 48 hours. The media was replaced with media containing 10 μg/mL (w/v) blasticidin, no puromycin or M3814 and incubated for 24 hours. For single cloning, approximately 500 cells were plated on a 10 cm dish in tissue culture media and allowed to grow for approximately one week. Single colonies were isolated and transferred to a 96 well plate using a microscope and 20 μL pipette tips. Clones were passaged and phenotypically screened for overnight cell death following 10 nM dTAG^V^-1 treatment, indicating an absence of wildtype SARS1. Homozygous knockin was confirmed with PCR of genomic DNA using the NucleoSpin Tissue kit (Macherey-Nagel 740952) and Q5 Hot Start Hi-Fidelity DNA polymerase (NEB M0494S). FWD primer 5’-CCCAGGACTGCAAGAACTGAT -3’ and REV primer 5’ CTTGGCTTCCCTGCTGTTG -3’. To generate C33A^SARS^^1-FH: E6^ cells, which express MTH1-E6 at the AAVS1 genomic site, C333A^SARS^^1-FH^ cells were seeded in 12 well plates at 6×10^6^ cells per well and incubated overnight. The following day, 0.4 μg of pKM5 and 0.6 μg of pKM6 were transfected with 1 μL PLUS reagent and 3 μL Lipofectamine LTX, delivered in 100 μL Opti-MEM. 6 hours after transfection, cells were passaged 1:6 into 6 well plates with 2.5 mL tissue culture medium and 1 μM M3814 and incubated for 24 hours. The media was replaced with media containing 1 μg/mL (w/v) puromycin and 1 μM M3814 and cells were incubated for 48 hours. The media was replaced with media containing 1 μg/mL (w/v) puromycin and no M3814 and incubated for 48 hours. The pool of cells was then expanded, cultured in media without puromycin, and used in subsequent assays.

### Chemical Induced Dimerization assay

For transient transfection assays, HEK293T cells were seeded for 24 hours in black clear-bottom 96 well plates (Corning 3904) at 2×10^4^ cells per well in 80 μL tissue culture media. Cells were then transfected with 90 ng total plasmid (60 ng ligase and 30 ng target), 0.1 μL PLUS reagent, and 0.25 μL Lipofectamine LTX, delivered in 10 μL of Opti-MEM and incubated for 24 hours. For experiments in engineered C33A cells, cells were plated in 90 μL of media and transfection was skipped. For inhibition assays, each well was treated with 0.5 μL of either DMSO or the noted inhibitors in 4.5 μL culture media for final concentrations of 10 μM MG132, 1 μM MLN4924, 5 μM AP1867, and 2 μM TH588 and treated for 15 minutes. Each well was then treated with 0.5 μL of a titration of FM4 in DMSO or DMSO only in 4.5 μL culture media to the noted final concentrations in a final well volume of 100 μL. Plates were then kept in the incubator for 24 hours. To quantify the abundance of HiBiT tagged target protein, 50 μL of Nano-Glo HiBiT Lytic Detection System (Promega N3040) was added to each well. Plates were shaken at 350 rpm for 10 minutes and incubated at room temperature for an additional 10 minutes before measurement with an EnVision plate reader. Luminescent values of treatment wells were divided by the DMSO-only wells of the same experimental conditions, then multiplied by 100 to find the percent target levels of control. A four-parameter non-linear regression curve fit was applied in GraphPad Prism to find EC_50_ concentrations, whereas D_max_ is reported as the average maximal degradation observed for each titration curve. For assays without additional chemical inhibitors or competitors, the same procedure was followed except that 0.5 μL of FM4 or dTAG^V^-1 titrations or DMSO only were added in 9.5 μL of culture media for a final well volume 100 μL. Different treatment times were used when indicated.

### Viability and apoptosis assays

The same cell seeding and compound dosing procedures as the Chemical Induced Dimerization assay were followed. To quantify cell viability 48 hours after treatment, plates were equilibrated to room temperature for 20 minutes and then 100 μL of room temperature CellTiter-Glo reagent (Promega G7570) was added to each well. Plates were shaken for 10 minutes at 350 rpm before measurement on an EnVision plate reader. To quantify apoptosis activity 6 hours after treatment, plates were equilibrated to room temperature and 100 μL of room temperature Caspase-Glo 3/7 reagent (Promega G8091) was added to each well. Plates were shaken for 10 minutes at 350 rpm and incubated at room temperature for an additional 20 minutes before measurement on an EnVison plate reader. Luminescent data were processed as described for the Chemical Induced Dimerization assay.

### Immunoprecipitation and immunoblotting

HEK293T cells were seeded in 12 well plates at 4×10^5^ cells per well transfected with 1 μg plasmid, 1.0 μL PLUS Reagent, and 2.5 μL Lipofectamine LTX, delivered in 100 μL Opti-MEM. After 24 hours, cells were washed with ice-cold PBS. HA-MTH1-vE3s immunoprecipitation was performed using Pierce Magnetic HA-Tag IP/Co-IP Kit (Thermo Scientfic 88838) as per manufacturer instructions, using 200 μL of lysis buffer plus Halt Protease Inhibitor (Thermo Scientific 78425). Input lysates and immunoprecipitated samples were separated by SDS-PAGE. Immunoblotting was done using rabbit-anti-HA (Cell Signaling 3724), rabbit-anti-DDB-1 (Cell Signaling 5428), rabbit-anti-E6AP (Cell Signaling 7526), and anti-rabbit-HRP (Cell Signaling 7074). C33A^SARS1-FH^ parental cells or cells stably expressing vE3s from the AAVS1 locus were seeded in 12 well plates at 6×10^5^ cells per well. After 24 hours, cells were washed with ice-cold PBS and lysed with RIPA Lysis and Extraction Buffer (Invitrogen 89900) plus cOmplete Protease Inhibitor Cocktail (Roche 04-693-116-001). Lysates were clarified by centrifugation at 21,000 x *g* for 10 minutes at 4 °C and separated by SDS-PAGE on 4-12% Bis-Tris polyacrylamide gel (Invitrogen NP0323BOX). Immunoblotting was done using rabbit-anti-GAPDH-HRP (Cell Signaling 3683), mouse-anti-p53 (Cell Signaling 2524), anti-mouse-HRP (Cytiva NA931).

### Knockdown effect on target degradation

For transient transfections in HEK293T, cells were seeded in black clear bottom 96 well plates at 1×10^4^ cells per well in 80 μL tissue culture media and incubated overnight. Cells were transfected with 1 pmol siRNA (Dharmacon ON-TARGETplus Smart Pools) and 0.2 μL Lipofectamine RNAiMAX, delivered in 5 μL Opti-MEM and incubated for 24 hours. Cells were then transfected with 90 ng total plasmid (60 ng ligase and 30 ng target), 0.1 μL PLUS Reagent, and 0.25 μL Lipofectamine LTX, delivered in 10 μL Opti-MEM and incubated for 24 hours. FM4 dissolved in DMSO or DMSO only was diluted into tissue culture media and added to wells, to a final FM4 concentration of 6 nM in 100 μL total volume. After 24 hours of treatment, HiBiT protein abundance was quantified as described in the Chemical Induced Dimerization assay section. To measure the extent of knockdown, HEK293T cells were seeded in 12 well plates at 1×10^5^ cells per well and transfected with 10 pmol siRNA and 2 μL Lipofectamine RNAiMAX. After incubation for 72 hours, cells were lysed with RIPA buffer, separated by SDS-PAGE, and immunoblotted as described in the immunoprecipitation and immunoblotting section. For C33A^SARS1-FH:E6^, cells were seeded in 6 well plates at 2.5×10^5^ cells per well in 2.2 mL tissue culture media and incubated overnight. Cells were transfected with 30 pmol siRNA and 7 μL Lipofectamine RNAiMAX, delivered in 300 μL Opti-MEM and incubated for 48 hours. 2.5 μL FM4 dissolved in DMSO or DMSO only was added directly to wells to a final concentration of 25 nM FM4, gently swirled to mix, and incubated for 24 hours. Cells were washed with PBS and lysed with 400 μL RIPA buffer plus cOmplete Protease Inhibitor Cocktail and clarified by centrifugation for at 21,000 x *g* for 10 minutes at 4°C. 5 μL of clarified lysate was added to 100 μL of room temperature Nano-Glo HibiT Lytic Detection System reagent or CellTiter-Glo reagent and quantitated by EnVision plate reader. HiBiT luminescence values were divided by CTG luminescence values for normalization, and then FM4 treated samples were divided by DMSO control samples and multiplied by100 to generate SARS1-FH levels as a percent of control. Clarified lysates were also separated by SDS-PAGE on 4-12% Bis-Tris polyacrylamide gel (Invitrogen NP0323BOX) and immunoblotted with rabbit-anti-SARS1 (Abcam ab154825), rabbit-anti-E6AP (Cell Signaling 7526), and rabbit-anti-GAPDH-HRP (Cell Signaling 3683),

### Metabolic labeling by puromoycin incorporation

C33A^SARS1-FH:E6^ cells were seeded in 12 well plates at 5×10^5^ cells per well and incubated overnight. Cells were treated with 30 μg/mL (w/v) cycloheximide (CHX), 10 nM dTAG^V^-1, 25 nM FM4, or a DMSO only control for 1 to 6 hr before adding 50 μg/mL (w/v) final concentration of puromycin for 30 minutes. Cells were washed with ice-cold PBS and lysed with RIPA Lysis and Extraction Buffer (Invitrogen 89900) plus cOmplete Protease Inhibitor Cocktail (Roche 04-693-116-001). Lysates were clarified by centrifugation at 21,000 x *g* for 10 minutes at 4 °C and separated by SDS-PAGE on 4-20% Tris-Glycine polyacrylamide gel (Invitrogen XP04205BOX). Immunoblotting was done using mouse-anti-puromycin (Sigma-Aldrich MABE343), rabbit-anti-GAPH (Cell Signaling 2118, anti-mouse-HRP (Cytiva NA931), and anti-rabbit AlexaFluor Plus 800 (Invitrogen A32735).

---

**Figure S1.**
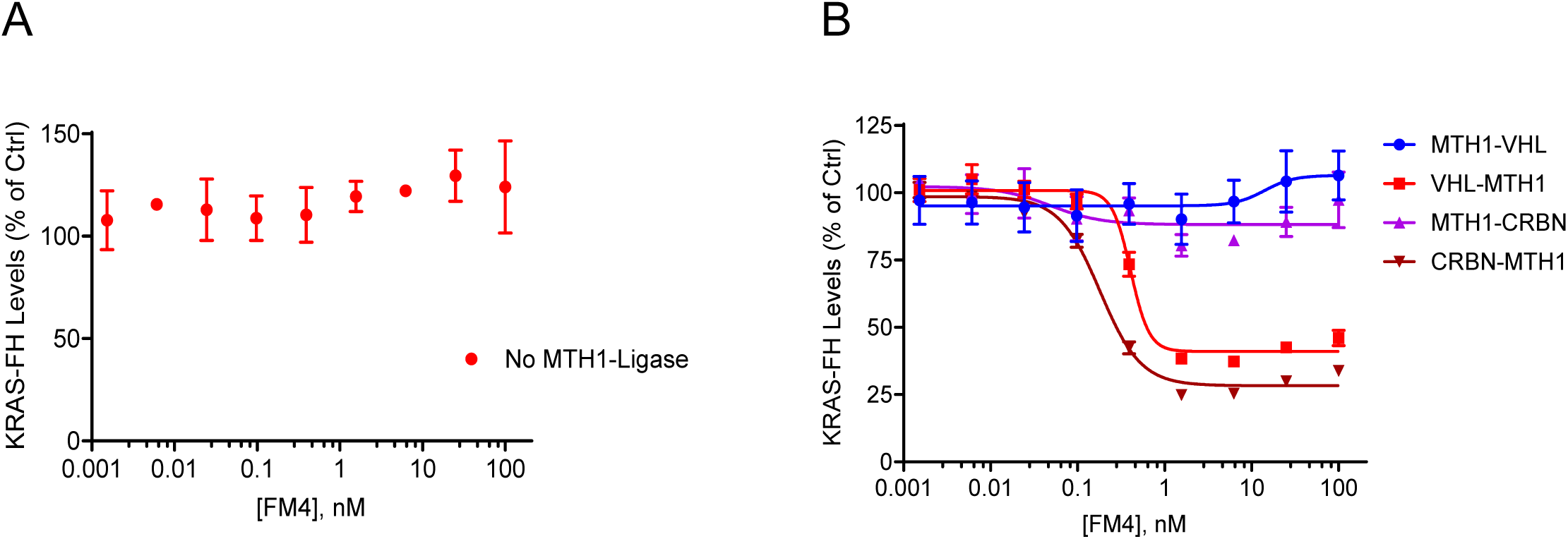
Target stability and ligase tag orientation in the chemical induced dimerization assay, related to Figure 1. A) FM4 treatment without MTH1 tagged ligase does not cause KRAS destabilization after 24 hours. B) C-terminal MTH1 tag orientation is superior in KRAS degradation with 24 hr FM4 treatment. All data representative of two independent experiments reported as the mean ± S.E.

**Figure S2.**
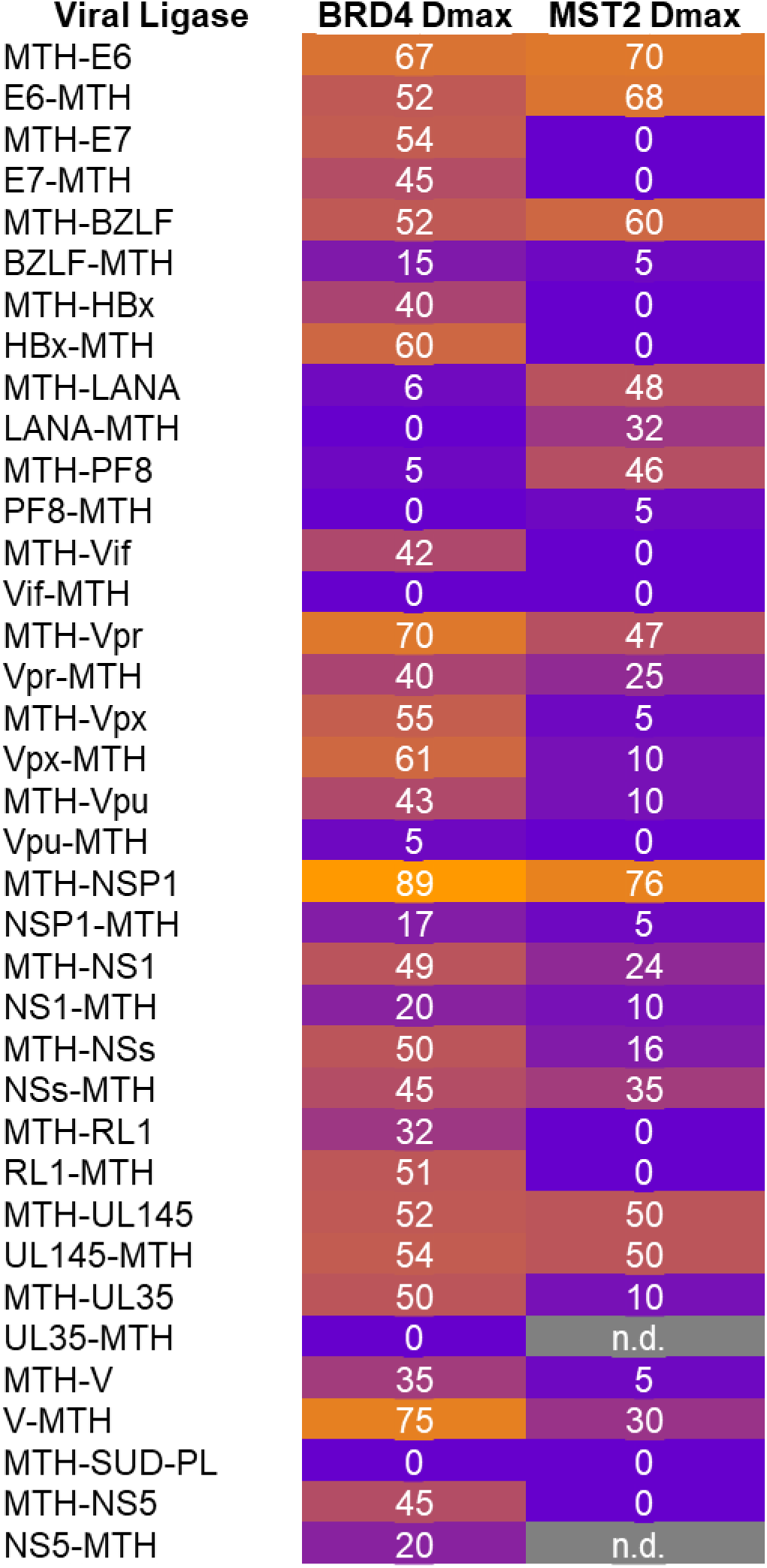
Effect of tag orientation on vE3 activity in CID assay, related to Figure 2. Dmax values from FM4 dose responses with N and C terminal MTH1 tag orientations of vE3s. Data representative of one experiment.

**Figure S3.**
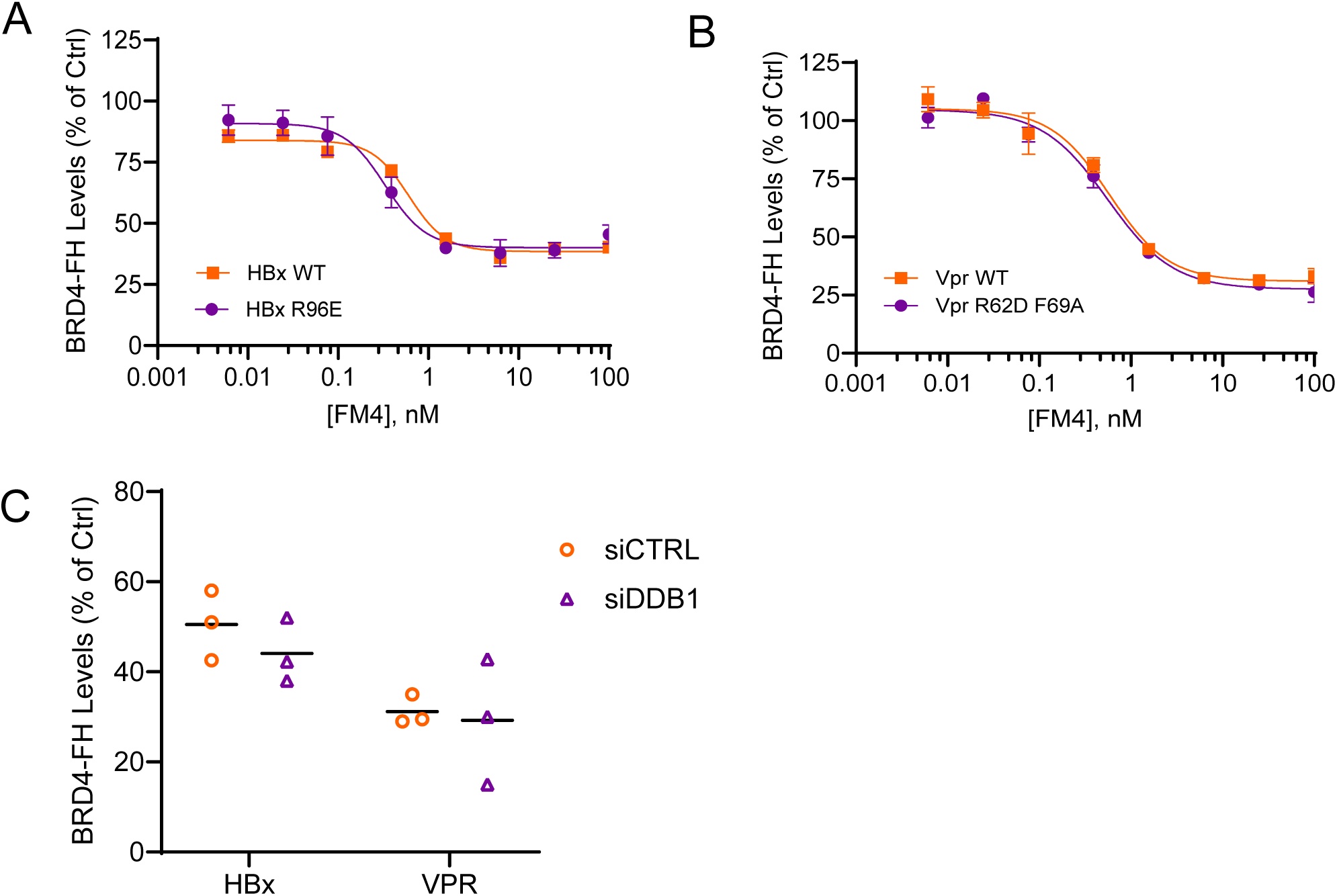
The vE3s HBx and Vpr degrade BRD4 through an unknown mechanism, related to Figure 3. A) HBx R96E mutation known to disrupt DDB1 binding does not reduce BRD4 degradation in CID assay. Data representative of two independent experiments reported as the mean ± S.E. B) Vpr double mutant known to disrupt DCAF1 binding does not reduce BRD4 degradation in CID assay. Data representative of two independent experiments reported as the mean ± S.E. C) HiBiT-BRD4^BD1/2^-FKBP12^F36V^ levels 24hr after treatment with 6 nM FM4 shows no change in degradation in DDB1 knockdown vs control cells. Individual data points from three independent experiments are presented, lines represent the means.

**Figure S4.**
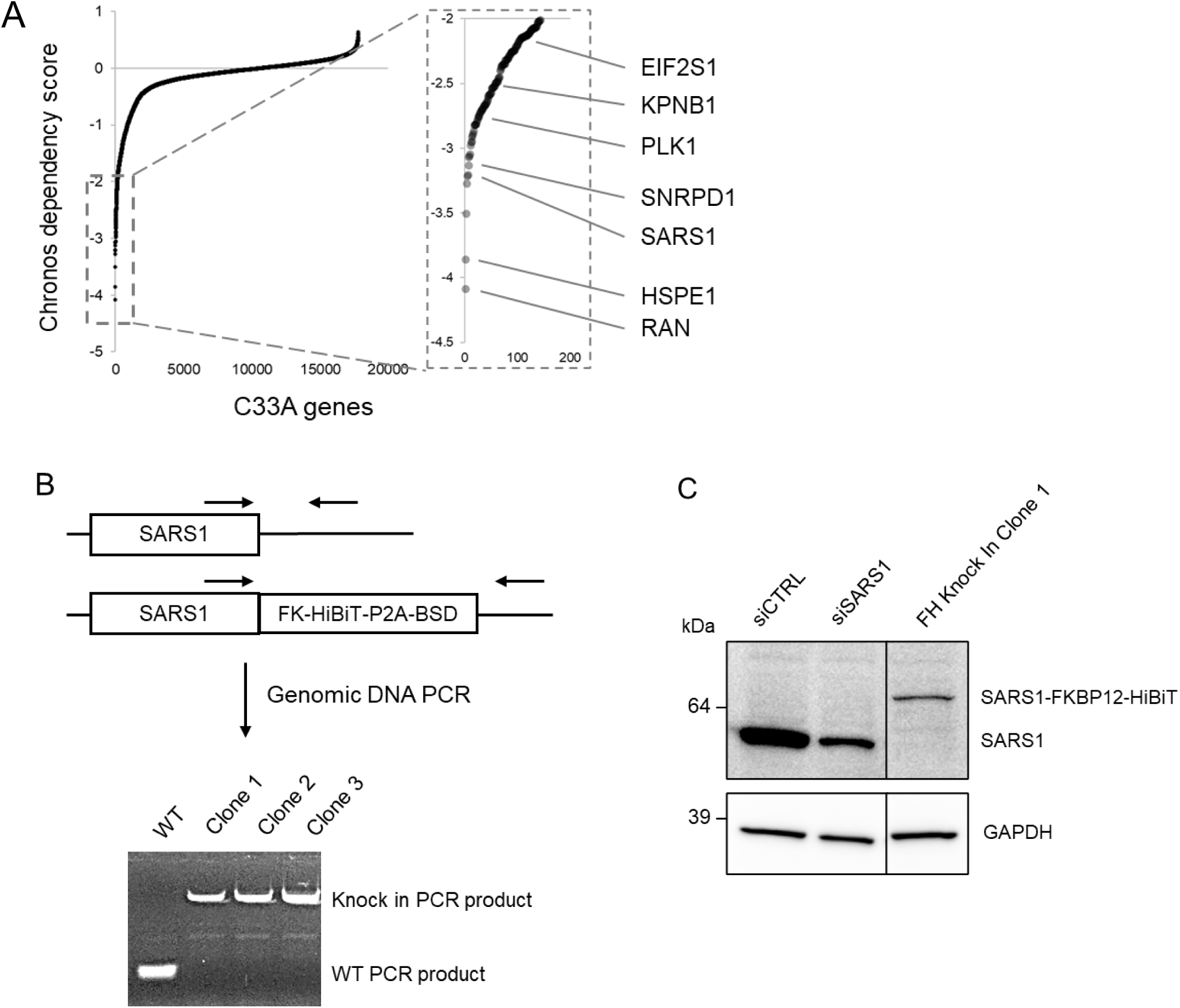
SARS1 knock-in in C33A cells, related to Figure 4. A) C33A genes ranked by Chronos dependency score. Insert shows genes below -2, with the genes tested in the CID assay marked. B) Genomic PCR of C33A SARS1-FH clones shows homozygous knock in at C-terminus of FKBP12(F36V)-HiBiT followed by the P2A cleavage sequence and blasticidin resistance gene, BSD. C) C33A SARS1-FH homozygous knock-in confirmed by Western blotting with anti-SARS1 antibody.

**Scheme-1:**
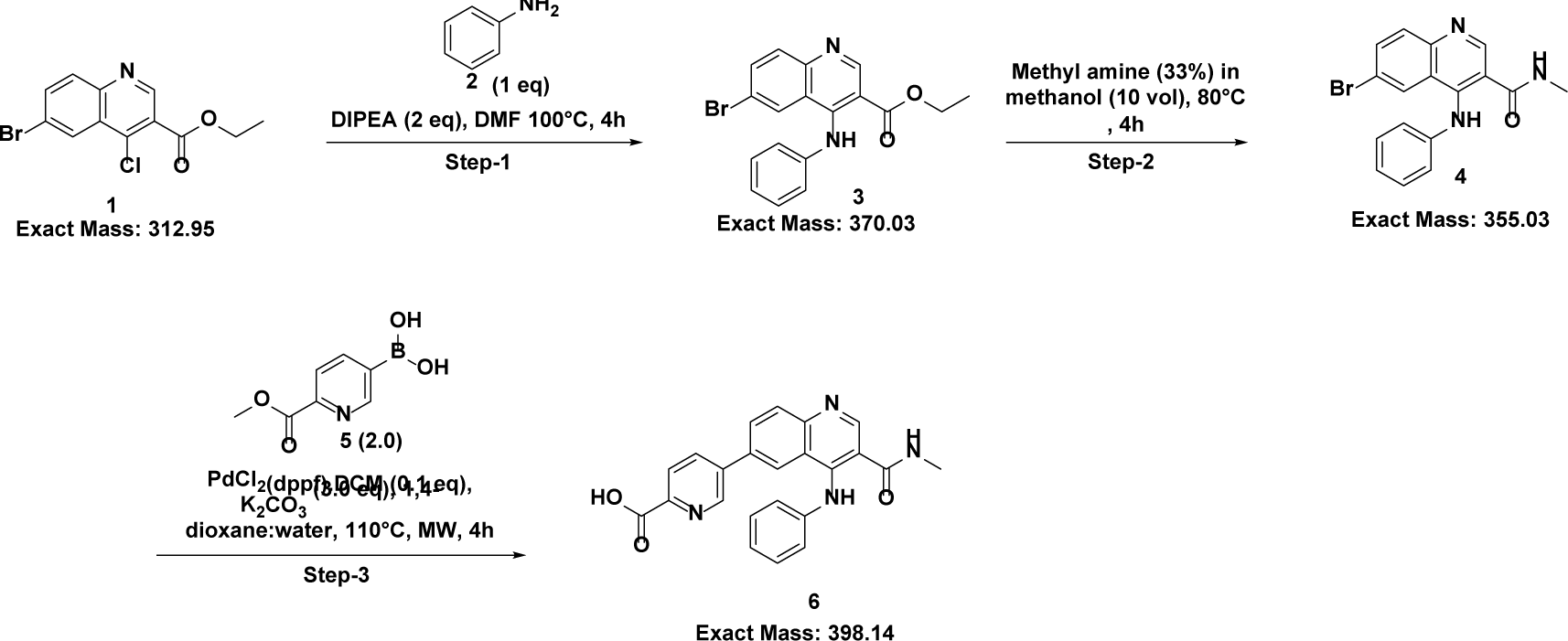
Synthesis of Inetermediate-6

**Scheme-2: Detailed.**
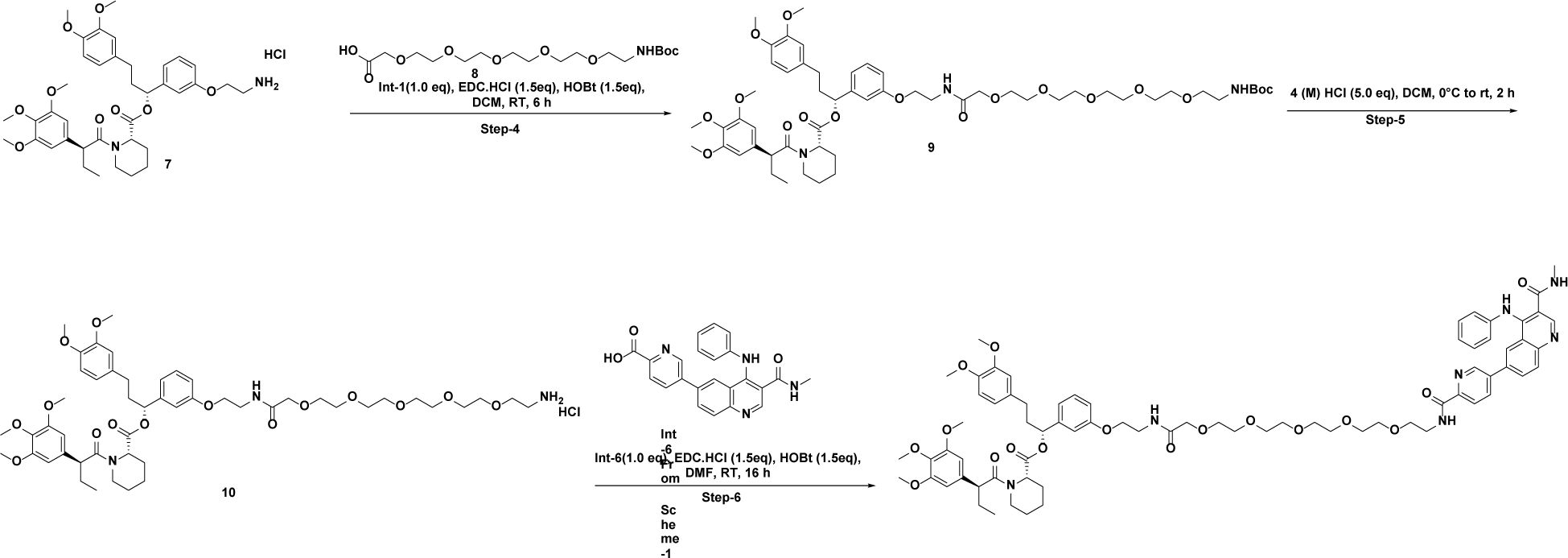
sytnesis routes and analytical data of FM4:

## CHEMICAL SYNTHESIS

### Synthesis schemes of FM4

### Step-1: Synthesis of ethyl 6-bromo-4-(phenylamino) quinoline-3-carboxylate

To a stirred solution of ethyl 6-bromo-4-chloroquinoline-3-carboxylate (5.0 g, 15.90 mmol) and aniline (1.451 mL, 15.90 mmol) in N, N-dimethylformamide (50.0 mL), was added DIPEA (5.55 mL, 31.8 mmol) at room temperature. The reaction mixture was stirred at 100 °C for 4h. After completion, the reaction mass poured into ice cold water (150 mL) and isolated the solid by filtration and dried, to give crude ethyl 6-bromo-4-(phenylamino) quinoline-3-carboxylate (5.0 g, 13.47 mmol, 85 % yield) as pale yellow solid.

### MS (ESI, positive ion) *m/z*: 372 (M+1)

**^1^H NMR (400 MHz, DMSO-*d*_6_)** δ 9.73 (s, 1H), 8.90 (s, 1H), 8.34 (dd, *J* = 1.8, 0.9 Hz, 1H), 7.98 – 7.80 (m, 2H), 7.33 (dd, *J* = 8.8, 7.0 Hz, 2H), 7.10 (m, 3H), 3.93 (q, *J* = 7.1 Hz, 2H), 1.12 (t, *J* = 7.1 Hz, 3H).

### Step-2: Synthesis of 6-bromo-N-methyl-4-(phenylamino) quinoline-3-carboxamide

Ethyl 6-bromo-4-(phenylamino) quinoline-3-carboxylate (5.2 g, 14.01 mmol) and methanamine in methanol (33%) (55.4 mL, 1401 mmol) were added in to a pressure vessel stirred at 80 °C for 3h. After completion, the reaction mixture was concentrated and the crude was purified by column chromatography through a Redi-Sep pre-packed silica gel column (80 g), eluting with a gradient of 80% to 90% EtOAc in hexane, to obtain 6-bromo-N-methyl-4-(phenylamino) quinoline-3-carboxamide (1.8 g, 5.05 mmol, 36.1 % yield) as a light-yellow solid.

### MS (ESI, positive ion) *m/z*: 356 (M+1)

**^1^H NMR (400 MHz, DMSO-*d*_6_)** δ 9.82 (s, 1H), 8.79 (s, 1H), 8.46 (d, *J* = 4.7 Hz, 1H), 8.24 (d, *J* = 2.0 Hz, 1H), 7.98 – 7.74 (m, 2H), 7.28 (dd, *J* = 7.0, 1.4 Hz, 1H), 7.08 – 6.96 (m, 2H), 2.52 (s, 3H).

### Step-3: Synthesis of 5-(3-(methylcarbamoyl)-4-(phenylamino) quinolin-6-yl) picolinic acid

A 20 mL glass microwave reaction vessel was charged with 6-bromo-N-methyl-4-(phenylamino) quinoline-3-carboxamide (300 mg, 0.842 mmol) and (6-(methoxycarbonyl) pyridin-3-yl) boronic acid (305 mg, 1.684 mmol), K_2_CO_3_ (349 mg, 2.53 mmol) in 1,4-dioxane (15 mL) and water (5.0 mL). The reaction mixture was degassed and PdCl_2_(dppf).CH_2_Cl_2_ was added. The reaction mixture was stirred in an Initiator microwave reactor at 110°C for 4h. The reaction mixture was diluted with water (10mL) and extracted with EtOAc (1 x 30 mL). The aqueous layer pH adjusted to 4-5 by using citric acid, and isolated the solid by filtration and dried, to give 5-(3-(methylcarbamoyl)-4-(phenylamino) quinolin-6-yl) picolinic acid (500 mg) as pale yellow solid.

Note: Two batches of 300 mg each were performed parallelly and filtered together.

### MS (ESI, positive ion) *m/z*: 399 (M+1)

**^1^H NMR (400 MHz, DMSO-*d*_6_)** δ 13.20 (s, 1H), 10.07 (s, 1H), 8.97 – 8.68 (m, 2H), 8.51 (d, *J* = 5.0 Hz, 1H), 8.39 (d, *J* = 2.1 Hz, 1H), 8.18 (ddd, *J* = 10.8, 8.4, 2.2 Hz, 2H), 8.08 (dd, *J* = 15.0, 8.4 Hz, 2H), 7.32 (t, *J* = 7.7 Hz, 2H), 7.17 – 7.01 (m, 3H), 2.56 (d, *J* = 4.5 Hz, 3H).

### Step-4: Synthesis of (R)-3-(3,4-dimethoxyphenyl)-1-(3-((2,2-dimethyl-4,22-dioxo-3,8,11,14,17,20-hexaoxa-5,23-diazapentacosan-25-yl)oxy)phenyl)propyl(S)-1-((S)-2-(3,4,5-trimethoyphe nyl)butanoyl)piperidine-2-carboxylate

To a stirred solution of **compound-7** (200 mg, 0.280 mmol,1.0 equiv) and 2,2-dimethyl-4-oxo-3,8,11,14-tetraoxa-5-azahexadecan-16-oic acid (86 mg, 0.280 mmol, 1.0 equiv) in dichloromethane (5 mL) was added EDC (80 mg, 0.419 mmol, 1.5 equiv), HOBt (64.2 mg, 0.419 mmol, 1.5 equiv) and DIPEA (244 μL, 1.398 mmol, 5.0 equiv) at 0 °C and stirred at RT for 6 h. After completion, the reaction mixture was diluted with water (20 mL) and extracted with CH_2_Cl_2_ (2 x 15 mL). The organic extract was washed with water (15 mL) and dried over Na_2_SO_4_. The solution was filtered and concentrated in vacuum to give the crude material as an orange oil. The crude material was absorbed onto a plug of silica gel and purified by column chromatography through a Redi-Sep pre-packed silica gel column (40 g), eluting with a gradient of 10% to 15% MeOH in DCM to provide **compound-9** (180 mg, 0.186 mmol, 66.5 % yield) as colorless thick oil.

**TLC solvent system:** 10% Methanol/DCM. Product’s R_f_: 0.6

MS (ESI, positive ion) *m/z*: 1056.4 [M+1]

### Step-5: Synthesis of (R)-1-(3-((20-amino-4-oxo-6,9,12,15,18-pentaoxa-3-**azaicosyl)oxy)phenyl)-3-(3,4-dimethoxyphenyl)propyl (S)-1-((S)-2-(3,4,5-trimethoxyphenyl)butanoyl)piperidine-2-carboxylate hydrochloride:**

A solution of **compound-9** (180 mg, 0.170 mmol, 1.0 equiv) in dichloromethane (5 mL) was added hydrogen chloride (4M in 1,4-Dioxane) (0.43 mL, 1.704 mmol, 10.0 equiv) at 0 °C and stirred at RT for 2 h. The reaction mixture was concentrated in vacuum to give the crude material as an orange oil. The crude material was washed with MTBE to provide **compound-10** (130 mg, 76.9 % yield) as an orange waxy solid.

**TLC solvent system:** 10% Methanol in DCM. Product’s R_f_: 0.1

**MS (ESI, positive ion) *m/z*:** 956.5 [M+1]

### Step-6: Synthesis of tert-butyl (R)-3-(3,4-dimethoxyphenyl)-1-(3-((1-(5-(3-(methylcarbamoyl)-4-(phenylamino)quinolin-6-yl)pyridin-2-yl)-1,19-dioxo-5,8,11,14,17-pentaoxa-2,20-diazadocosan-22-yl)oxy)phenyl)propyl (S)-1-((S)-2-(3,4,5-trimethoxyphenyl)butanoyl)piperidine-2-carboxylate

To a solution of **compound-10** (149 mg, 0.151 mmol, 1.0 equiv) and 5-(3-(methylcarbamoyl)-4-(phenylamino)quinolin-6-yl)picolinic acid (60 mg, 0.151 mmol, 1.0 equiv) in N, N-dimethylformamide (10 mL) was added EDC (43.3 mg, 0.226 mmol, 1.5 equiv), HOBt (34.6 mg, 0.226 mmol, 1.5 equiv) and DIPEA (0.13 mL, 0.755 mmol, 5 equiv) at 0 °C and stirred at RT for 6 h. Progress of the reaction was monitored by UPLC. The reaction mixture was diluted with water (15 mL) and extracted with EtOAc (3 x15 mL). The organic extract was dried over Na_2_SO_4_. The solution was filtered and concentrated under vacuum to give the crude material as a orange oil. The crude material was purified by reverse-phase preparative HPLC using a Kinetex C8 (250 x 21.2)mm 5.0µm, 0.1% FA in water B:-ACN to FM4 (65 mg, 0.049 mmol, 32.3 % yield) as yellow solid.

**TLC solvent system:** 10% Methanol in DCM. Product’s R_f_: 0.6

**MS (ESI, positive ion) *m/z*:** 1336.5 [M+1]

**^1^H NMR (400 MHz, DMSO-*d_6_*)** δ 10.07 (s, 1H), 8.83 (s, 1H), 8.77 (dd, *J* = 2.3, 0.8 Hz, 1H), 8.71 (t, *J* = 5.8 Hz, 1H), 8.52 (d, *J* = 4.7 Hz, 1H), 8.37 (d, J = 2.0 Hz, 1H), 8.18 (ddd, *J* = 16.1, 8.5, 2.2 Hz, 2H), 8.10 (d, *J* = 0.9 Hz, 1H), 8.08 (d, *J* = 1.5 Hz, 1H), 7.84 (t, *J* = 5.9 Hz, 1H), 7.31 (dd, *J* = 8.3, 7.3 Hz, 3H), 7.15 (t, J = 7.9 Hz, 1H), 7.10 – 7.00 (m, 3H), 6.91 (s, 1H), 6.82 (d, *J* = 8.2 Hz, 2H), 6.78 (s, 1H), 6.72 (d, *J* = 2.0 Hz, 1H), 6.64 (d, *J* = 1.9 Hz, 1H), 6.62 (s, 1H), 6.56 (s, 1H), 6.54 (s, 1H), 5.52 (dd, *J* = 8.4, 5.1 Hz, 1H), 5.27 (s, 1H), 4.00 (t, *J* = 5.8 Hz, 3H), 3.89 (s, 2H), 3.86 (s, 1H), 3.74 (s, 2H), 3.73 – 3.68 (m, 6H), 3.64 (s, 1H), 3.59 (s, 4H), 3.57 (s, 3H), 3.56 – 3.51 (m, 10H), 3.49 (d, *J* = 2.6 Hz, 11H), 2.57 (d, *J* = 4.6 Hz, 4H), 2.15 (s, 2H), 1.99 – 1.76 (m, 3H), 1.58 (dd, *J* = 18.1, 11.6 Hz, 5H), 1.24 (s, 1H), 0.81 (t, *J* = 7.2 Hz, 3H).

## REFERENCES

1. Gao, H., Sun, X., and Rao, Y. (2020). PROTAC Technology: Opportunities and Challenges. ACS Med Chem Lett 11, 237–240. 10.1021/acsmedchemlett.9b00597.

2. Verma, R., Mohl, D., and Deshaies, R.J. (2020). Harnessing the Power of Proteolysis for Targeted Protein Inactivation. Mol Cell 77, 446–460. 10.1016/j.molcel.2020.01.010.

3. Samarasinghe, K.T.G., and Crews, C.M. (2021). Targeted protein degradation: A promise for undruggable proteins. Cell Chem Biol 28, 934–951. 10.1016/j.chembiol.2021.04.011.

4. Bondeson, D.P., Smith, B.E., Burslem, G.M., Buhimschi, A.D., Hines, J., Jaime-Figueroa, S., Wang, J., Hamman, B.D., Ishchenko, A., and Crews, C.M. (2018). Lessons in PROTAC Design from Selective Degradation with a Promiscuous Warhead. Cell Chem Biol 25, 78–87 e75. 10.1016/j.chembiol.2017.09.010.

5. Huang, H.T., Dobrovolsky, D., Paulk, J., Yang, G., Weisberg, E.L., Doctor, Z.M., Buckley, D.L., Cho, J.H., Ko, E., Jang, J., et al. (2018). A Chemoproteomic Approach to Query the Degradable Kinome Using a Multi-kinase Degrader. Cell Chem Biol 25, 88–99 e86. 10.1016/j.chembiol.2017.10.005.

6. Testa, A., Lucas, X., Castro, G.V., Chan, K.H., Wright, J.E., Runcie, A.C., Gadd, M.S., Harrison, W.T.A., Ko, E.J., Fletcher, D., and Ciulli, A. (2018). 3-Fluoro-4-hydroxyprolines: Synthesis, Conformational Analysis, and Stereoselective Recognition by the VHL E3 Ubiquitin Ligase for Targeted Protein Degradation. J Am Chem Soc 140, 9299–9313. 10.1021/jacs.8b05807.

7. Han, X., Zhao, L., Xiang, W., Qin, C., Miao, B., Xu, T., Wang, M., Yang, C.Y., Chinnaswamy, K., Stuckey, J., and Wang, S. (2019). Discovery of Highly Potent and Efficient PROTAC Degraders of Androgen Receptor (AR) by Employing Weak Binding Affinity VHL E3 Ligase Ligands. J Med Chem 62, 11218–11231. 10.1021/acs.jmedchem.9b01393.

8. Donovan, K.A., Ferguson, F.M., Bushman, J.W., Eleuteri, N.A., Bhunia, D., Ryu, S., Tan, L., Shi, K., Yue, H., Liu, X., et al. (2020). Mapping the Degradable Kinome Provides a Resource for Expedited Degrader Development. Cell 183, 1714–1731 e1710. 10.1016/j.cell.2020.10.038.

9. Liu, Y., Yang, J., Wang, T., Luo, M., Chen, Y., Chen, C., Ronai, Z., Zhou, Y., Ruppin, E., and Han, L. (2023). Expanding PROTACtable genome universe of E3 ligases. Nat Commun 14, 6509. 10.1038/s41467-023-42233-2.

10. Weng, G., Cai, X., Cao, D., Du, H., Shen, C., Deng, Y., He, Q., Yang, B., Li, D., and Hou, T. (2023). PROTAC-DB 2.0: an updated database of PROTACs. Nucleic Acids Res 51, D1367–D1372. 10.1093/nar/gkac946.

11. Houzelstein, D., Bullock, S.L., Lynch, D.E., Grigorieva, E.F., Wilson, V.A., and Beddington, R.S. (2002). Growth and early postimplantation defects in mice deficient for the bromodomain-containing protein Brd4. Mol Cell Biol 22, 3794–3802. 10.1128/MCB.22.11.3794-3802.2002.

12. Winter, G.E., Buckley, D.L., Paulk, J., Roberts, J.M., Souza, A., Dhe-Paganon, S., and Bradner, J.E. (2015). DRUG DEVELOPMENT. Phthalimide conjugation as a strategy for in vivo target protein degradation. Science 348, 1376–1381. 10.1126/science.aab1433.

13. Guenette, R.G., Yang, S.W., Min, J., Pei, B., and Potts, P.R. (2022). Target and tissue selectivity of PROTAC degraders. Chem Soc Rev 51, 5740–5756. 10.1039/d2cs00200k.

14. Khan, S., Zhang, X., Lv, D., Zhang, Q., He, Y., Zhang, P., Liu, X., Thummuri, D., Yuan, Y., Wiegand, J.S., et al. (2019). A selective BCL-X(L) PROTAC degrader achieves safe and potent antitumor activity. Nat Med 25, 1938–1947. 10.1038/s41591-019-0668-z.

15. Donovan, K.A., An, J., Nowak, R.P., Yuan, J.C., Fink, E.C., Berry, B.C., Ebert, B.L., and Fischer, E.S. (2018). Thalidomide promotes degradation of SALL4, a transcription factor implicated in Duane Radial Ray syndrome. Elife 7. 10.7554/eLife.38430.

16. Surka, C., Jin, L., Mbong, N., Lu, C.C., Jang, I.S., Rychak, E., Mendy, D., Clayton, T., Tindall, E., Hsu, C., et al. (2021). CC-90009, a novel cereblon E3 ligase modulator, targets acute myeloid leukemia blasts and leukemia stem cells. Blood 137, 661–677. 10.1182/blood.2020008676.

17. Mahon, C., Krogan, N.J., Craik, C.S., and Pick, E. (2014). Cullin E3 ligases and their rewiring by viral factors. Biomolecules 4, 897–930. 10.3390/biom4040897.

18. Dybas, J.M., Herrmann, C., and Weitzman, M.D. (2018). Ubiquitination at the interface of tumor viruses and DNA damage responses. Curr Opin Virol 32, 40–47. 10.1016/j.coviro.2018.08.017.

19. Liu, Y., and Tan, X. (2020). Viral Manipulations of the Cullin-RING Ubiquitin Ligases. Adv Exp Med Biol 1217, 99–110. 10.1007/978-981-15-1025-0_7.

20. Scheffner, M., Huibregtse, J.M., Vierstra, R.D., and Howley, P.M. (1993). The HPV-16 E6 and E6-AP complex functions as a ubiquitin-protein ligase in the ubiquitination of p53. Cell 75, 495–505. 10.1016/0092-8674(93)90384-3.

21. Zanier, K., Charbonnier, S., Sidi, A.O., McEwen, A.G., Ferrario, M.G., Poussin-Courmontagne, P., Cura, V., Brimer, N., Babah, K.O., Ansari, T., et al. (2013). Structural basis for hijacking of cellular LxxLL motifs by papillomavirus E6 oncoproteins. Science 339, 694–698. 10.1126/science.1229934.

22. Martinez-Zapien, D., Ruiz, F.X., Poirson, J., Mitschler, A., Ramirez, J., Forster, A., Cousido-Siah, A., Masson, M., Vande Pol, S., Podjarny, A., et al. (2016). Structure of the E6/E6AP/p53 complex required for HPV-mediated degradation of p53. Nature 529, 541–545. 10.1038/nature16481.

23. Wang, J.C.K., Baddock, H.T., Mafi, A., Foe, I.T., Bratkowski, M., Lin, T.Y., Jensvold, Z.D., Preciado Lopez, M., Stokoe, D., Eaton, D., et al. (2024). Structure of the p53 degradation complex from HPV16. Nat Commun 15, 1842. 10.1038/s41467-024-45920-w.

24. Li, T., Robert, E.I., van Breugel, P.C., Strubin, M., and Zheng, N. (2010). A promiscuous alpha-helical motif anchors viral hijackers and substrate receptors to the CUL4-DDB1 ubiquitin ligase machinery. Nat Struct Mol Biol 17, 105–111. 10.1038/nsmb.1719.

25. Le-Trilling, V.T.K., Becker, T., Nachshon, A., Stern-Ginossar, N., Scholer, L., Voigt, S., Hengel, H., and Trilling, M. (2020). The Human Cytomegalovirus pUL145 Isoforms Act as Viral DDB1-Cullin-Associated Factors to Instruct Host Protein Degradation to Impede Innate Immunity. Cell Rep 30, 2248–2260 e2245. 10.1016/j.celrep.2020.01.070.

26. Wick, E.T., Treadway, C.J., Li, Z., Nicely, N.I., Ren, Z., Baldwin, A.S., Xiong, Y., Harrison, J.S., and Brown, N.G. (2022). Insight into Viral Hijacking of CRL4 Ubiquitin Ligase through Structural Analysis of the pUL145-DDB1 Complex. J Virol 96, e0082622. 10.1128/jvi.00826-22.

27. Mui, U.N., Haley, C.T., and Tyring, S.K. (2017). Viral Oncology: Molecular Biology and Pathogenesis. J Clin Med 6. 10.3390/jcm6120111.

28. Tempera, I., and Lieberman, P.M. (2021). Oncogenic Viruses as Entropic Drivers of Cancer Evolution. Front Virol 1. 10.3389/fviro.2021.753366.

29. Benhenda, S., Cougot, D., Buendia, M.A., and Neuveut, C. (2009). Hepatitis B virus X protein molecular functions and its role in virus life cycle and pathogenesis. Adv Cancer Res 103, 75–109. 10.1016/S0065-230X(09)03004-8.

30. Diao, M.K., Liu, C.Y., Liu, H.W., Li, J.T., Li, F., Mehryar, M.M., Wang, Y.J., Zhan, S.B., Zhou, Y.B., Zhong, R.G., and Zeng, Y. (2015). Integrated HPV genomes tend to integrate in gene desert areas in the CaSki, HeLa, and SiHa cervical cancer cell lines. Life Sci 127, 46–52. 10.1016/j.lfs.2015.01.039.

31. Levrero, M., and Zucman-Rossi, J. (2016). Mechanisms of HBV-induced hepatocellular carcinoma. J Hepatol 64, S84–S101. 10.1016/j.jhep.2016.02.021.

32. Hoppe-Seyler, K., Bossler, F., Braun, J.A., Herrmann, A.L., and Hoppe-Seyler, F. (2018). The HPV E6/E7 Oncogenes: Key Factors for Viral Carcinogenesis and Therapeutic Targets. Trends Microbiol 26, 158–168. 10.1016/j.tim.2017.07.007.

33. Sivasudhan, E., Blake, N., Lu, Z., Meng, J., and Rong, R. (2022). Hepatitis B Viral Protein HBx and the Molecular Mechanisms Modulating the Hallmarks of Hepatocellular Carcinoma: A Comprehensive Review. Cells 11. 10.3390/cells11040741.

34. Montrose, K., and Krissansen, G.W. (2014). Design of a PROTAC that antagonizes and destroys the cancer-forming X-protein of the hepatitis B virus. Biochem Biophys Res Commun 453, 735–740. 10.1016/j.bbrc.2014.10.006.

35. de Wispelaere, M., Du, G., Donovan, K.A., Zhang, T., Eleuteri, N.A., Yuan, J.C., Kalabathula, J., Nowak, R.P., Fischer, E.S., Gray, N.S., and Yang, P.L. (2019). Small molecule degraders of the hepatitis C virus protease reduce susceptibility to resistance mutations. Nat Commun 10, 3468. 10.1038/s41467-019-11429-w.

36. Desantis, J., Mercorelli, B., Celegato, M., Croci, F., Bazzacco, A., Baroni, M., Siragusa, L., Cruciani, G., Loregian, A., and Goracci, L. (2021). Indomethacin-based PROTACs as pan-coronavirus antiviral agents. Eur J Med Chem 226, 113814. 10.1016/j.ejmech.2021.113814.

37. Xu, Z., Liu, X., Ma, X., Zou, W., Chen, Q., Chen, F., Deng, X., Liang, J., Dong, C., Lan, K., et al. (2022). Discovery of oseltamivir-based novel PROTACs as degraders targeting neuraminidase to combat H1N1 influenza virus. Cell Insight 1, 100030. 10.1016/j.cellin.2022.100030.

38. Alugubelli, Y.R., Xiao, J., Khatua, K., Kumar, S., Sun, L., Ma, Y., Ma, X.R., Vulupala, V.R., Atla, S., Blankenship, L.R., et al. (2024). Discovery of First-in-Class PROTAC Degraders of SARS-CoV-2 Main Protease. J Med Chem 67, 6495–6507. 10.1021/acs.jmedchem.3c02416.

39. Emert-Sedlak, L.A., Tice, C.M., Shi, H., Alvarado, J.J., Shu, S.T., Reitz, A.B., and Smithgall, T.E. (2024). PROTAC-mediated degradation of HIV-1 Nef efficiently restores cell-surface CD4 and MHC-I expression and blocks HIV-1 replication. Cell Chem Biol 31, 658–668 e614. 10.1016/j.chembiol.2024.02.004.

40. Grifagni, D., Lenci, E., De Santis, A., Orsetti, A., Barracchia, C.G., Tedesco, F., Bellini Puglielli, R., Lucarelli, F., Lauriola, A., Assfalg, M., et al. (2024). Development of a GC-376 Based Peptidomimetic PROTAC as a Degrader of 3-Chymotrypsin-like Protease of SARS-CoV-2. ACS Med Chem Lett 15, 250–257. 10.1021/acsmedchemlett.3c00498.

41. Liu, H.Y., Li, Z., Reindl, T., He, Z., Qiu, X., Golden, R.P., Donovan, K.A., Bailey, A., Fischer, E.S., Zhang, T., et al. (2024). Broad-spectrum activity against mosquito-borne flaviviruses achieved by a targeted protein degradation mechanism. Nat Commun 15, 5179. 10.1038/s41467-024-49161-9.

42. Zhao, J., Wang, J., Pang, X., Liu, Z., Li, Q., Yi, D., Zhang, Y., Fang, X., Zhang, T., Zhou, R., et al. (2022). An anti-influenza A virus microbial metabolite acts by degrading viral endonuclease PA. Nat Commun 13, 2079. 10.1038/s41467-022-29690-x.

43. Raina, K., Forbes, C.D., Stronk, R., Rappi, J.P., Jr., Eastman, K.J., Gerritz, S.W., Yu, X., Li, H., Bhardwaj, A., Forgione, M., et al. (2023). Regulated Induced Proximity Targeting Chimeras (RIPTACs): a Novel Heterobifunctional Small Molecule Therapeutic Strategy for Killing Cancer Cells Selectively. bioRxiv. 10.1101/2023.01.01.522436.

44. Nabet, B., Roberts, J.M., Buckley, D.L., Paulk, J., Dastjerdi, S., Yang, A., Leggett, A.L., Erb, M.A., Lawlor, M.A., Souza, A., et al. (2018). The dTAG system for immediate and target-specific protein degradation. Nat Chem Biol 14, 431–441. 10.1038/s41589-018-0021-8.

45. Veits, G.K., Henderson, C.S., Vogelaar, A., Eron, S.J., Lee, L., Hart, A., Deibler, R.W., Baddour, J., Elam, W.A., Agafonov, R.V., et al. (2021). Development of an AchillesTAG degradation system and its application to control CAR-T activity. Current Research in Chemical Biology 1. 10.1016/j.crchbi.2021.100010.

46. Clackson, T., Yang, W., Rozamus, L.W., Hatada, M., Amara, J.F., Rollins, C.T., Stevenson, L.F., Magari, S.R., Wood, S.A., Courage, N.L., et al. (1998). Redesigning an FKBP-ligand interface to generate chemical dimerizers with novel specificity. Proc Natl Acad Sci U S A 95, 10437–10442. 10.1073/pnas.95.18.10437.

47. Gad, H., Koolmeister, T., Jemth, A.S., Eshtad, S., Jacques, S.A., Strom, C.E., Svensson, L.M., Schultz, N., Lundback, T., Einarsdottir, B.O., et al. (2014). MTH1 inhibition eradicates cancer by preventing sanitation of the dNTP pool. Nature 508, 215–221. 10.1038/nature13181.

48. Salsman, J., Jagannathan, M., Paladino, P., Chan, P.K., Dellaire, G., Raught, B., and Frappier, L. (2012). Proteomic profiling of the human cytomegalovirus UL35 gene products reveals a role for UL35 in the DNA repair response. J Virol 86, 806–820. 10.1128/JVI.05442-11.

49. Grant, A., Ponia, S.S., Tripathi, S., Balasubramaniam, V., Miorin, L., Sourisseau, M., Schwarz, M.C., Sanchez-Seco, M.P., Evans, M.J., Best, S.M., and Garcia-Sastre, A. (2016). Zika Virus Targets Human STAT2 to Inhibit Type I Interferon Signaling. Cell Host Microbe 19, 882–890. 10.1016/j.chom.2016.05.009.

50. Ma-Lauer, Y., Carbajo-Lozoya, J., Hein, M.Y., Muller, M.A., Deng, W., Lei, J., Meyer, B., Kusov, Y., von Brunn, B., Bairad, D.R., et al. (2016). p53 down-regulates SARS coronavirus replication and is targeted by the SARS-unique domain and PLpro via E3 ubiquitin ligase RCHY1. Proc Natl Acad Sci U S A 113, E5192–5201. 10.1073/pnas.1603435113.

51. Nightingale, K., Potts, M., Hunter, L.M., Fielding, C.A., Zerbe, C.M., Fletcher-Etherington, A., Nobre, L., Wang, E.C.Y., Strang, B.L., Houghton, J.W., et al. (2022). Human cytomegalovirus protein RL1 degrades the antiviral factor SLFN11 via recruitment of the CRL4 E3 ubiquitin ligase complex. Proc Natl Acad Sci U S A 119. 10.1073/pnas.2108173119.

52. Parisien, J.P., Lenoir, J.J., Alvarado, G., and Horvath, C.M. (2022). The Human STAT2 Coiled-Coil Domain Contains a Degron for Zika Virus Interferon Evasion. J Virol 96, e0130121. 10.1128/JVI.01301-21.

53. Ramakrishnan, D., Xing, W., Beran, R.K., Chemuru, S., Rohrs, H., Niedziela-Majka, A., Marchand, B., Mehra, U., Zabransky, A., Dolezal, M., et al. (2019). Hepatitis B Virus X Protein Function Requires Zinc Binding. J Virol 93. 10.1128/JVI.00250-19.

54. Fabryova, H., and Strebel, K. (2019). Vpr and Its Cellular Interaction Partners: R We There Yet? Cells 8. 10.3390/cells8111310.

55. de Martel, C., Georges, D., Bray, F., Ferlay, J., and Clifford, G.M. (2020). Global burden of cancer attributable to infections in 2018: a worldwide incidence analysis. Lancet Glob Health 8, e180–e190. 10.1016/S2214-109X(19)30488-7.

56. Shin, H.J., Kim, J.Y., Hampson, L., Pyo, H., Baek, H.J., Roberts, S.A., Hendry, J.H., and Hampson, I.N. (2010). Human papillomavirus 16 E6 increases the radiosensitivity of p53-mutated cervical cancer cells, associated with up-regulation of aurora A. Int J Radiat Biol 86, 769–779. 10.3109/09553002.2010.484477.

57. Lu, Y., Chen, Y., Zhang, Z., Li, M., Chen, X., Tu, K., and Li, L. (2022). HPV16 E6 promotes cell proliferation, migration, and invasion of human cervical cancer cells by elevating both EMT and stemness characteristics. Cell Biol Int 46, 599–610. 10.1002/cbin.11756.

58. Schmidt, E.K., Clavarino, G., Ceppi, M., and Pierre, P. (2009). SUnSET, a nonradioactive method to monitor protein synthesis. Nat Methods 6, 275–277. 10.1038/nmeth.1314.

59. Schiller, J.T., Castellsague, X., and Garland, S.M. (2012). A review of clinical trials of human papillomavirus prophylactic vaccines. Vaccine 30 *Suppl 5*, F123–138. 10.1016/j.vaccine.2012.04.108.

60. Cohen, P.A., Jhingran, A., Oaknin, A., and Denny, L. (2019). Cervical cancer. Lancet 393, 169–182. 10.1016/S0140-6736(18)32470-X.

61. Chitsike, L., and Duerksen-Hughes, P.J. (2021). PPI Modulators of E6 as Potential Targeted Therapeutics for Cervical Cancer: Progress and Challenges in Targeting E6. Molecules 26. 10.3390/molecules26103004.

62. Messa, L., and Loregian, A. (2022). HPV-induced cancers: preclinical therapeutic advancements. Expert Opin Investig Drugs 31, 79–93. 10.1080/13543784.2021.2010703.

63. Ye, X., Zhang, P., Tao, J., Wang, J.C.K., Mafi, A., Grob, N.M., Quartararo, A.J., Baddock, H.T., Chan, L.J.G., McAllister, F.E., et al. (2023). Discovery of reactive peptide inhibitors of human papillomavirus oncoprotein E6. Chem Sci 14, 12484–12497. 10.1039/d3sc02782a.

64. Wang, X., Sandberg, M.L., Martin, A.D., Negri, K.R., Gabrelow, G.B., Nampe, D.P., Wu, M.L., McElvain, M.E., Toledo Warshaviak, D., Lee, W.H., et al. (2021). Potent, Selective CARs as Potential T-Cell Therapeutics for HPV-positive Cancers. J Immunother 44, 292–306. 10.1097/CJI.0000000000000386.

65. Stachura, P., Stencel, O., Lu, Z., Borkhardt, A., and Pandyra, A.A. (2023). Arenaviruses: Old viruses present new solutions for cancer therapy. Front Immunol 14, 1110522. 10.3389/fimmu.2023.1110522.

66. Morreale, F.E., Kleine, S., Leodolter, J., Junker, S., Hoi, D.M., Ovchinnikov, S., Okun, A., Kley, J., Kurzbauer, R., Junk, L., et al. (2022). BacPROTACs mediate targeted protein degradation in bacteria. Cell 185, 2338–2353 e2318. 10.1016/j.cell.2022.05.009.

67. Hoi, D.M., Junker, S., Junk, L., Schwechel, K., Fischel, K., Podlesainski, D., Hawkins, P.M.E., van Geelen, L., Kaschani, F., Leodolter, J., et al. (2023). Clp-targeting BacPROTACs impair mycobacterial proteostasis and survival. Cell 186, 2176–2192 e2122. 10.1016/j.cell.2023.04.009.

68. Lee, A.K., and Potts, P.R. (2017). A Comprehensive Guide to the MAGE Family of Ubiquitin Ligases. J Mol Biol 429, 1114–1142. 10.1016/j.jmb.2017.03.005.

